# Genetic slippage after sex maintains diversity for parasite resistance in a natural host population

**DOI:** 10.1101/2021.07.11.451958

**Authors:** Camille Ameline, Felix Vögtli, Jason Andras, Eric Dexter, Jan Engelstädter, Dieter Ebert

**Affiliations:** Department of Environmental Sciences, Zoology, University of Basel, Vesalgasse 1, 4051 Basel, Switzerland; Department of Biological Sciences, Clapp Laboratory, Mount Holyoke College, South Hadley, MA, USA; School of Biological Sciences, The University of Queensland, Brisbane, Australia

**Keywords:** cost of sex, parasite-mediated selection, zooplankton, resistance, resting egg, resting stage, sex, dormancy, genetic model of resistance, *Daphnia magna*, *Pasteuria ramosa*

## Abstract

Although parasite-mediated selection is thought to be a major driver of host evolution, its influence on genetic variation for parasite resistance is not yet well understood. We monitored a large population of the planktonic crustacean *Daphnia magna* over eight years, as it underwent yearly epidemics of the bacterial pathogen *Pasteuria ramosa*. We observed a cyclical pattern of resistance evolution: resistant phenotypes increased in frequency throughout the epidemics, but susceptibility was restored each spring when hosts hatched from sexual resting stages, a phenomenon described as genetic slippage in response to sex. Collecting and hatching *D. magna* resting stages across multiple seasons showed that largely resistant host populations can produce susceptible offspring through recombination. Resting stages produced throughout the planktonic season accurately represent the hatching population cohort of the following spring. A genetic model of resistance developed for this host–parasite system, based on multiple loci and strong epistasis, is in partial agreement with these findings. Our results reveal that, despite strong selection for resistance in a natural host population, genetic slippage after sexual reproduction has the potential to maintain genetic diversity of host resistance.

## Introduction

The origin and maintenance of diversity is a major question in evolutionary biology, with the respective roles of selection, mutation, and drift in maintaining genetic diversity in nature still being disputed (Fisher 1930; Fisher 1958; Frank 2013; Kern and Hahn 2018). Parasites (including pathogens) have been suggested as a causal factor for some highly diverse regions in plant and animal genomes. Indeed, the role of selection by parasites is well established for the major histocompatibility (MHC) gene complex in jawed vertebrates and resistance (R) genes in plants (Jeffery and Bangham 2000; Hughes 2002; Radwan et al. 2020), both of which have remarkably high allelic diversity (Sommer 2005; Baggs et al. 2017). In particular, coevolution with parasites is linked to increased host diversity (Wang et al. 2017; Duxbury et al. 2019), and high diversity at resistance genes has been shown to be advantageous against parasites (Sommer 2005; Zhao et al. 2016; Gösser et al. 2019; Peters et al. 2019; White et al. 2020).

As a key mechanism for creating diversity via novel allele combinations, sexual reproduction is a central component of host-parasite coevolution theory (Lively 2010; Morran et al. 2011; Auld et al. 2016). Recombination may allow a host population to create new genotypes that parasites are not yet adapted to, thereby escaping parasites adapted to specific host genotypes. Based on this reasoning, it has been suggested that parasites select for the maintenance of host sexual reproduction as a mechanism to create and maintain beneficial genetic diversity—the Red Queen theory (Jaenike 1978; Bell and Smith 1987; Hamilton et al. 1990; MacPherson and Otto 2018). Coevolution with parasites has indeed been shown to promote sex and outbreeding (Morran et al. 2011; Gibson et al. 2016), and there is empirical evidence of the advantage of sexual over asexual reproduction in natural systems and associated experiments (Tobler and Schlupp 2008; Auld et al. 2016; Gibson et al. 2018). The Red Queen theory posits that parasite interactions make sexual recombination advantageous for hosts as they can produce and benefit from rare allele combinations. Selection should then have the form of time-lagged negative frequency-dependent selection (NFDS) (Hamilton et al. 1990; Salathé et al. 2008; Lively 2010). On the other hand, sexual reproduction may represent a cost for a population that has evolved adaptive resistance to a parasite because it may destroy advantageous allele combinations (Otto and Nuismer 2004; Otto 2009). Models have shown that recombination could be selected against under certain conditions of genetic architecture and selection strength (Engelstädter and Bonhoeffer 2009; Kouyos et al. 2009; Engelstädter 2015), resulting in distinct dynamics of resistance and infectivity genotypes and phenotypes in hosts and parasites populations.

Coevolution by NFDS can maintain genetic diversity through balancing selection within and among populations (reviewed in Ebert and Fields 2020). Red Queen dynamics assume specific forms of host–parasite interactions without which polymorphisms at loci under selection would soon disappear (Agrawal and Lively 2002; Otto and Nuismer 2004; Thrall et al. 2016). A major assumption of the specific genetic interaction matrices of Red Queen models is epistasis and—for diploid organisms—dominance. For most host-parasite systems, however, we know very little about the link between the genetic architecture of resistance, the effect of selection on resistance, and the role of genetic recombination for the evolution of the system. In this study we aim to fill this gap for a cyclic parthenogenetic crustacean that undergoes seasonal epidemics of a virulent bacterial pathogen.

To understand the role of sexual recombination in shaping the evolutionary dynamics of resistance, knowledge of the genetic architecture of resistance is necessary. Empirical work determining resistance to parasites in natural systems has suggested a multi-locus architecture, with dominance and epistasis, for most systems (Sasaki 2000; Li and Cowling 2003; Tellier and Brown 2007; Wilfert and Schmid-Hempel 2008; Metzger et al. 2016). Dominance and epistasis (i.e. nonadditive gene action) are particularly important because they determine the degree to which the mean and variance of phenotypes change after genetic recombination (Otto and Nuismer 2004; Otto 2009). Depending on the form and strength of natural selection and the mode of gene action determining the trait under selection, recombination may impede or enhance the response to selection (Lynch and Deng 1994; Otto 2009).

Many organisms in diverse taxa, such as cladocerans, monogonont rotifers, bryozoan, and aphids, reproduce by cyclical parthenogenesis. They produce parthenogenetic eggs directly throughout most of the season, with occasional periods of sexual reproduction that result in resting stages that usually hatch at the beginning of the following season (Decaestecker et al. 2009). In such a reproductive system, selection is expected to increase the mean fitness of the population. After a round of sexual recombination, the mean fitness of the population presumably decreases, a phenomenon known as regression to the mean before selection (Falconer 1981), or genetic slippage in response to sex (Lynch and Deng 1994; Decaestecker et al. 2009). The variance of the trait under selection is also expected to change, although the direction of the change cannot be easily predicted as it depends on the signs of the covariances between genetic effects in the parental generation (Lynch and Deng 1994). In rotifer populations, variance has been observed to both increase and decrease after sexual reproduction (Becks and Agrawal 2011; Becks and Agrawal 2012). Due to their extended period of asexual reproduction, cyclic parthenogens are good systems to study genetic slippage (Becks and Agrawal 2011). During the asexual phase, selection over time can build up stronger linkage disequilibria among loci and deviations from Hardy-Weinberg equilibrium (Lynch and Deng 1994), making the effect of both selection and recombination on the trait under selection more evident.

We monitored resistance phenotypic changes over eight consecutive years in a large natural population of the crustacean *Daphnia magna*, whose yearly population cycle includes strong summer epidemics of the bacterial parasite *Pasteuria ramosa,* sexual reproduction to survive the winter, and the hatching of sexual resting stages in spring. In a previous study, we documented parasite-mediated selection and resolved parts of the underlying genetic architecture of resistance to the local parasite (Ameline et al. 2021). Here we reveal genetic slippage created by sexual reproduction in this cyclical parthenogen, showing that resistance increases during the yearly parasite epidemics, but that recombination reestablishes the initial resistance diversity in the following planktonic season. We link this long-term monitoring to a system-specific genetic model of resistance that predicts the impact of sexual reproduction on the temporal dynamics of the evolution of resistance. Our data suggest that dominance and epistasis are crucial for explaining the maintenance of genetic diversity for resistance in this host population, but also reveal that our current understanding of the mechanisms at work does not account for the full complexity of the genetics of resistance.

## Results

### Seasonal epidemics

We monitored a large *Daphnia magna* population in the fishless pond Aegelsee, Switzerland (Ameline et al. 2021) from 09.10.2010 to 24.09.2018, observing strong annual epidemics of *Pasteuria ramosa* that typically started about a month after the host emerged from diapause in early April and lasted through most of the summer (Fig. 1A). Epidemics reached peak prevalence of 70% to nearly 100%; no epidemic of any other known *D. magna* parasite was observed in this population. The population overwinters exclusively in the form of sexually produced resting stages, with an estimated overwintering population size of several millions. We also monitored environmental and ecological variables over the course of our study and present those results in Supplementary Fig. S1.

**Figure 1.**
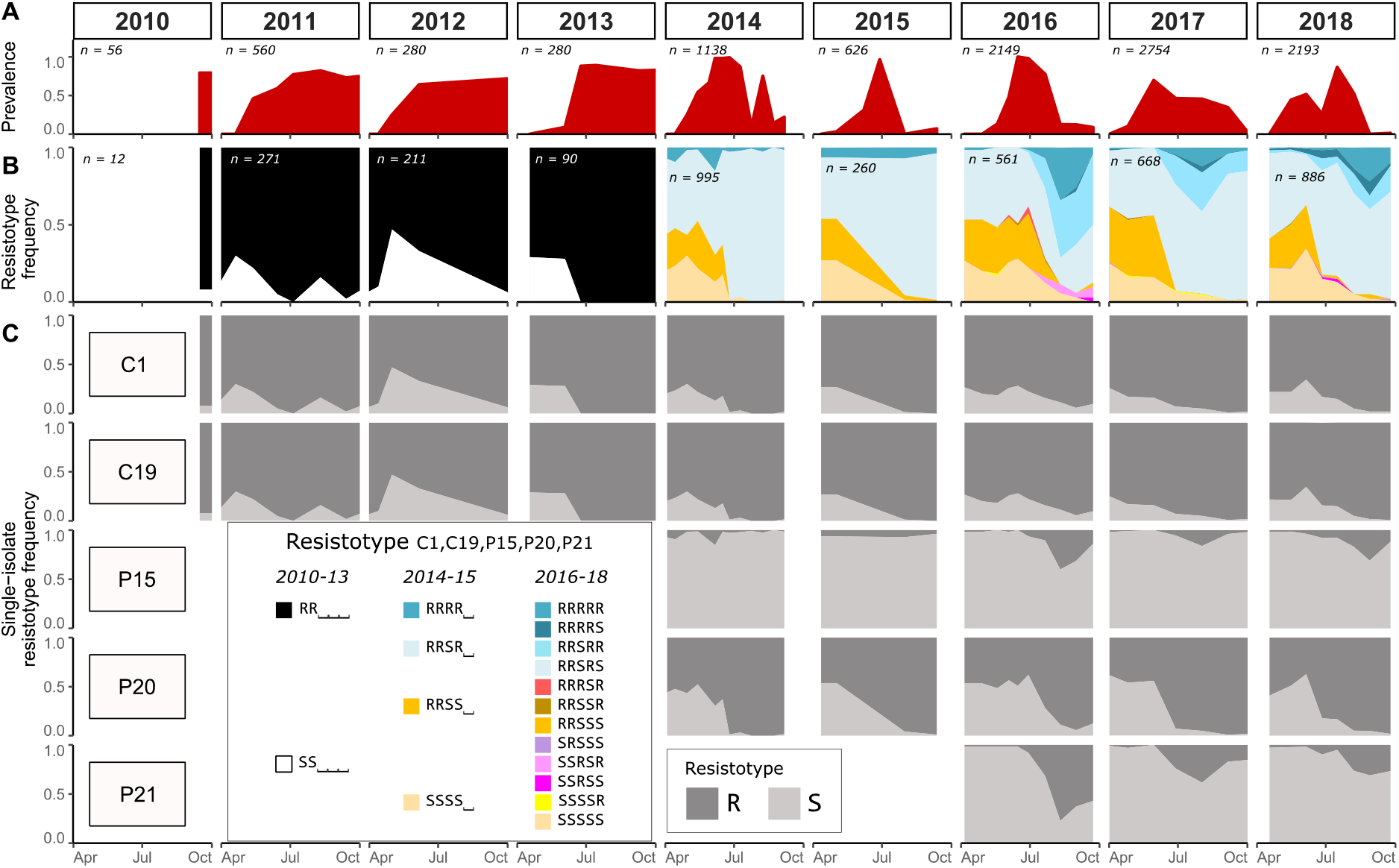
Cyclic resistotype dynamics across eight years in the Aegelsee. From 2010 to 2018, samples of *Daphnia magna* were collected from early April to early October every two to four weeks. Parasite prevalence was recorded and about 60 to 100 animals were cloned and their resistotypes (resistance phenotypes) assessed. **A**: *Pasteuria ramosa* prevalence (= proportion of infected females) in the *D. magna* population. **B**: Resistotype frequency in the *D. magna* population. Resistance and susceptibility to individual *P. ramosa* isolates are denoted as R and S, respectively. The combined resistotype shows resistance for up to five *P. ramosa* isolates: C1, C19, P15, P20 and P21. Until 2013 only C1 and C19 were tested; in 2014 and 2015 isolates C1, C19, P15 and P20 were tested; and all five isolates were tested after 2015. We use the placeholder “⎵”when an isolate was not tested. Resistance to P20 is pinpointed because of its importance in the evolution of the host population (Ameline et al. 2021). The “n” denotes the total number of genotypes tested in a given year. **C**: Resistotype frequency to each of the five *P. ramosa* isolates. Note the strong increase in resistance to P20 every year.

### Resistotype dynamics

Using five isolates of the pathogen *Pasteuria ramosa*, we quantified the proportion of resistant (R) and susceptible (S) host phenotypes in the population with an attachment test that measures the parasite’s ability to attach to the host cuticle; failure to attach indicates resistant hosts (Duneau et al. 2011). Because we can clone females using the host’s parthenogenetic eggs (iso-female lines), we can perform this test on several individuals with the same genotype. Resistotypes—i.e. resistance phenotypes—are presented as a sequence of R and S letters, each letter representing resistance or susceptibility to one of the five tested parasite isolates in the following order: C1, C19, P15, P20 and P21. We used the placeholder “⎵” for isolates that we did not test or consider.

For eight successive years we observed similar resistotype frequencies in spring (Fig. 1B). From 2011 to 2013, when data resolution was lower due to less frequent and smaller samples, the spring cohort was composed of about 25% of the SS⎵⎵⎵ resistotype and 75% of the RR⎵⎵⎵ resistotype (Fig. 1B). Two additional *P. ramosa* isolates were added from 2014 and one more from 2016 onwards, allowing for a more refined picture that was dominated by four phenotypes: resistotypes SSSS⎵ and RRSS⎵ each represented about 25% of the population; RRSR⎵ about 45%; and RRRR⎵ about 5% (Fig. 1B). Overall, R resistotypes dominated for C1, C19 and P20, while S resistotypes dominated for P15 and P21 (Fig. 1C).

Each year, these resistotype frequencies were relatively stable at the beginning of the season, but changed dramatically after the start of the *P. ramosa* epidemic in May. Two resistant phenotypes, namely RRSR⎵ and RRRR⎵ (blue in Fig. 1B, 2014-2018), increased in proportion, while the resistotypes susceptible to P15 and P20—RRSS⎵ and SSSS⎵—decreased in proportion (orange and yellow in Fig. 1B). Overall, resistance to all individual *P. ramosa* strains increased over the season (dark gray in Fig. 1C): resistance to C1 and C19 increased every year from 79±2% to 97±1% during the entire six months of the *D. magna* planktonic phase. The biggest change was resistance to P20, which increased from 49±4% to 96±2% within two months during the main peak of the epidemics. Resistance to P15 and P21 showed a more complex pattern, with a tendency to increase during the second half of the summer and decrease again towards the end of the season (Fig. 1C).

The stable spring frequencies across years, together with the strong dynamics across the summer season, resulted in a strong pattern of cyclic resistotype frequencies changes. Among about four thousand tested genotypes across eight years, some resistotypes were never observed in our samples, e.g. SS⎵R⎵ and RS⎵⎵⎵, possibly indicating genetically impossible phenotypes combinations or absence of polymorphisms at the underlying resistance loci in this population.

### Response to selection for resistance

As a cyclic parthenogen, *D. magna* reproduces asexually during most of the active season and produces sexual resting stages (embryos) in a protective case (ephippium) that overwinter and hatch in the spring. Because the planktonic animals do not overwinter in our population, the spring cohort is exclusively the result of sexual reproduction. To look at the impact of selection and recombination on resistance diversity during and between seasons, respectively, we calculated the mean and variance of resistance phenotypes of the planktonic population for each sample through time, assigning resistance (R) and susceptibility (S) a value of 1 and 0, respectively. If directional selection acts on resistance, we expect mean resistance to increase, as selection removes susceptible phenotypes. As hardly any susceptible resistotypes are left at the end of the summer, we further expected variance in resistance to decrease during the summer. Furthermore, a round of sexual reproduction is expected to restore, or partly restore, the variance and the mean is expected to relapse to some degree because genetic recombination leads (under most conditions) to a regress to the mean before selection (Falconer 1981)—also discussed as genetic slippage (Lynch and Deng 1994; Decaestecker et al. 2009). Our results align with these predictions: every year, mean resistance increased, and variance declined over the planktonic season (Fig. 2). After sexual reproduction, variance was restored, and the mean regressed towards the mean of the previous year before selection. What was surprising, however, was that the relapse of the mean was nearly perfect over the entire study period, indicating that there was no overall response to selection across seasons. Note that the apparent drop in the mean between 2015 and 2016 in Figure 2 is caused by the addition of one more *P. ramosa* isolate (P21) in the test panel.

**Figure 2.**
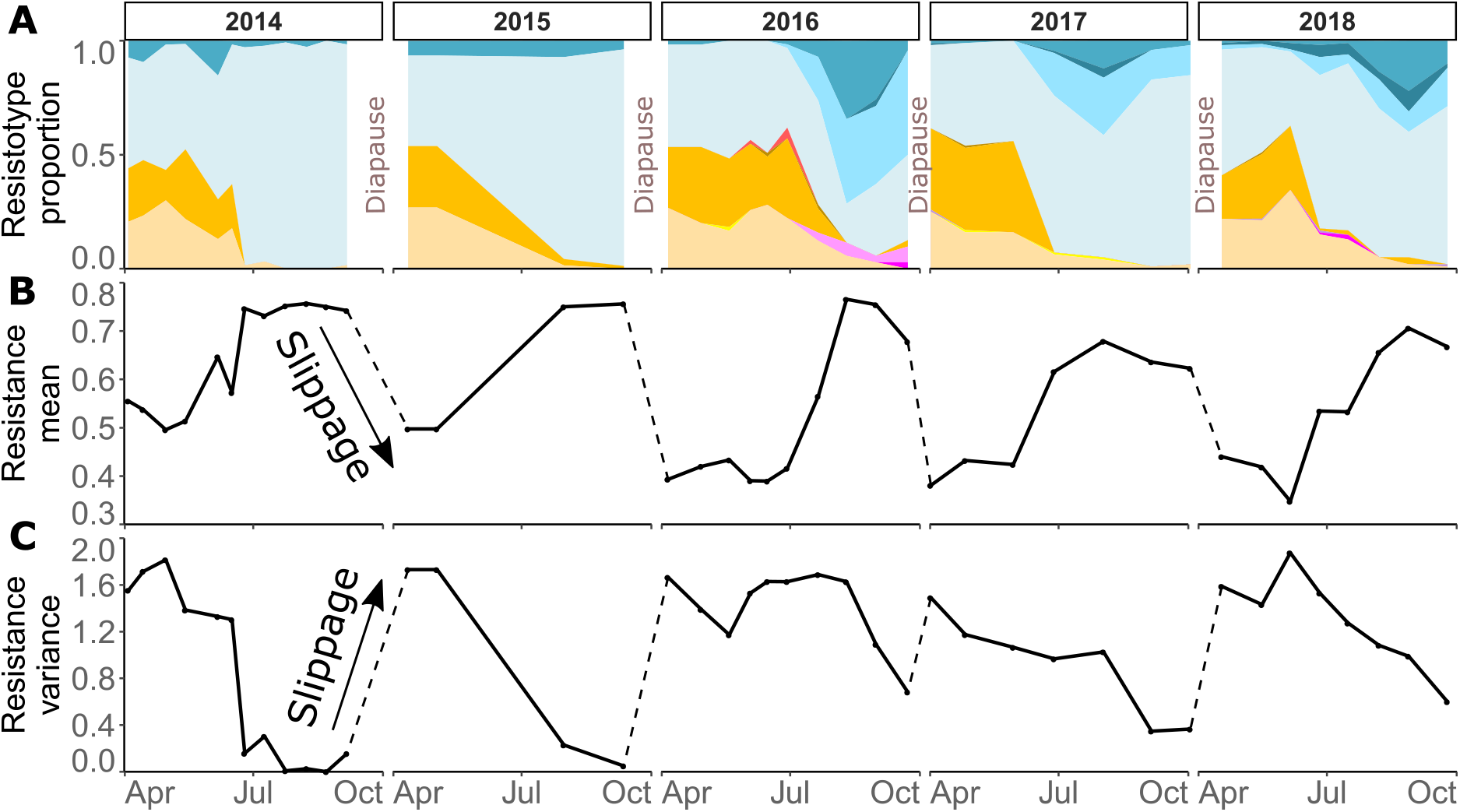
Genetic slippage resulting from sexual reproduction in the Aegelsee *Daphnia magna* population. **A**: Observed resistotype (resistance phenotype) frequencies in the *D. magna* population from 2014 to 2018 (same as Fig. 1B for 2014 to 2018; repeated here for better comparison). **B**: Mean resistance to *Pasteuria ramosa* across time. Mean resistance increases across every summer planktonic phase. We attributed to each resistotype a resistance score ranging from zero to the number of isolates tested, and weighted the mean per sampling point by the number of tested isolates, resulting in a score between zero and one (e.g. RRRRR would have an overall resistance score of 1 and SSSSS would be 0). The dashed lines span the time windows during which sexual offspring overwinter and hatch the following spring. **C**: Variance of resistance across time, calculated along with the mean in panel B. Note that in 2014 and 2015, four bacterial isolates were tested, while we used five from 2016 to 2018. Therefore, mean and variance cannot be directly compared between years when different number of parasite isolates are used.

### Selection and sexual reproduction

To understand these pronounced dynamics in mean resistance and its variance, we collected and hatched sexually produced resting stages across three seasons, using sediment traps that we emptied at about monthly intervals. Sediment traps allow us to decouple the current resting egg production from resting eggs produced earlier, as these traps only collect resting stages that are dropped from the current planktonic population. This allowed us to estimate when sexual reproduction occurred and—by subsequent hatching of resting stages from each sampling date—to estimate the hatchlings resistotype frequencies.

We observed that resting stages were produced throughout most of *D. magna*’s planktonic phase and tended to show multiple peaks before and after the main change in resistotype frequencies in June-July (Fig 3B). The number of resting stages per ephippium (zero, one or two) produced over the planktonic phase of *D. magna* remained approximately stable (Supplementary Fig. S2B and C, linear regression, all years pooled: *R*^*2*^ = 0.14, *F* = 3.2 on 1 and 13 DF, *p* = 0.095). After overwintering, the overall hatching success in outdoor containers was 74.4±3.9%, which was stable across the planktonic phase when the resting stages were produced (linear regression, pooled for all years: *R*^*2*^ = 0.042, *F* = 1.75 on 1 and 16 DF, *p* = 0.20). The hatching pattern after induction was also consistent, with most resting stages hatching within a few days after induction (Supplementary Fig. S2). The few resting stages that hatched later did not differ in their resistotype proportions from the earlier hatchlings (measured only in 2014, Fisher’s test, *p* = 0.32, Supplementary Fig. S3).

**Figure 3.**
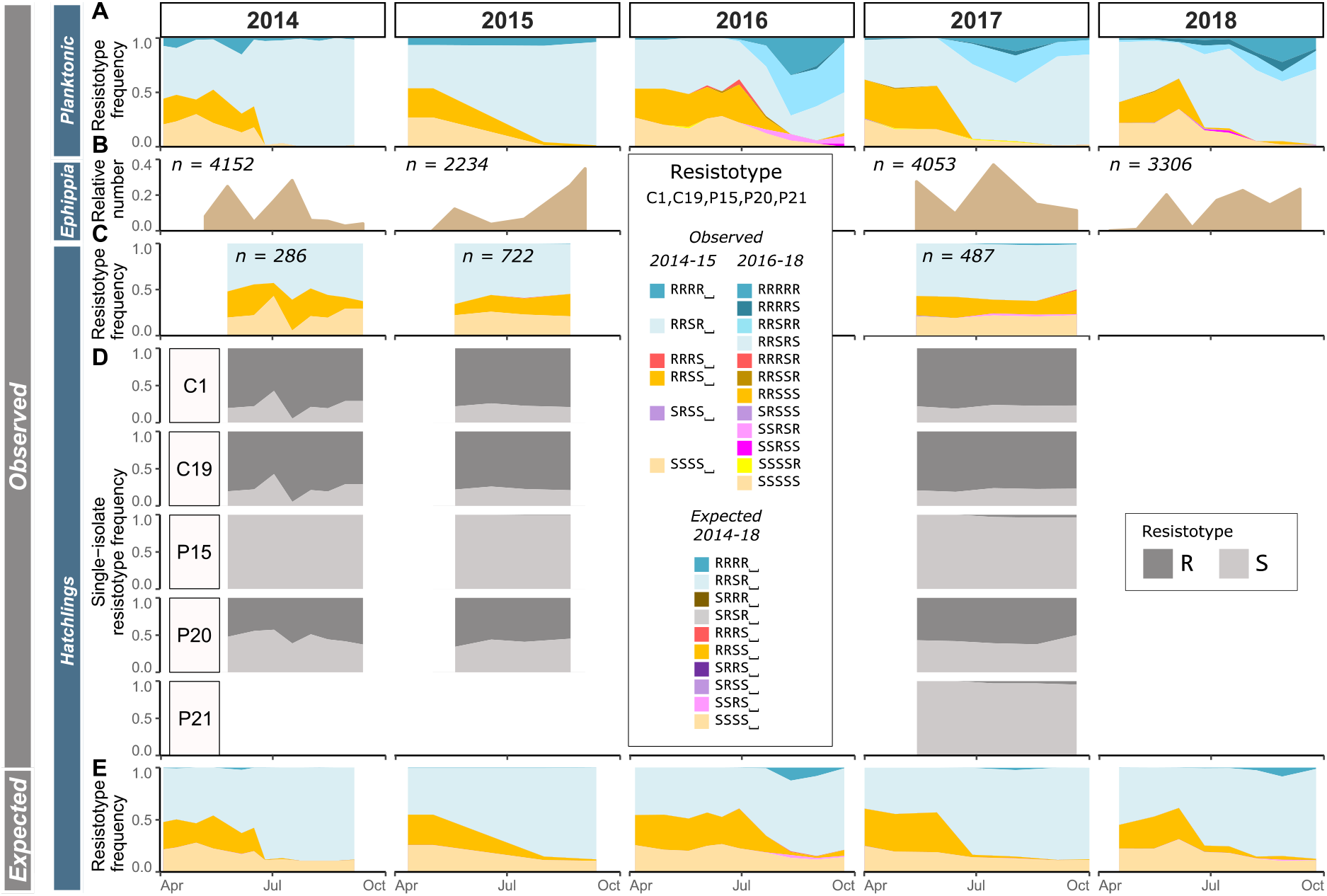
Longitudinal resting stage hatching of *Daphnia magna* from the Aegelsee. **A**: Observed resistotype (resistance phenotype) frequencies in the *D. magna* population from 2014 to 2018 (same as Fig. 1B, repeated here for better comparison). **B**: Observed relative number of *D. magna* resting stage cases (ephippia) produced in the pond and recovered from five to nine sediment traps, in two-four week intervals from early April to early October in 2014, 2015, 2017 and 2018. Time on the x-axis represents the mid-point between two consecutive emptying of the traps. The “n” indicates the total number of ephippia for a given year. **C**: Resistotype frequencies of the hatchlings from the sediment traps plotted against the collection time (only for 2014, 2015 and 2017). Resting stages from 2018 were collected but not hatched. Note that in 2014, the first resting stage sample was lost. In 2015, no hatchlings emerged from the last sample. We represent the four-letter resistotype (C1, C19, P15, P20) to be comparable with the E panel. **D**: As in **C** but for each of the five *Pasteuria ramosa* isolates separately. **E**: Expected resistotype frequencies of hatchlings from sexually produced eggs (resting stages) by the planktonic population across the entire planktonic season (also for parts of the season where no resting stages were produced). Expected resistotype frequencies were calculated using the genetic model of resistance in the *D. magna–P. ramosa* system assuming random mating of the parent population at the time of resting stage production. Detailed methods and results of these calculations are given in the text and in Supplementary Figs. S5 and S6 and Tables S1 and S2.

All hatchlings were cloned and tested for resistotypes. Surprisingly, in all years, the observed resistotype frequencies of the hatchlings remained rather stable over the season, both for the combined and for the individual bacterial isolates (Fig. 3C and D), independent of the strongly changing resistotype composition of the planktonic animals at the time of resting egg production (Fig. 3A). This created a substantial difference between the resistotype distribution of the parent population and their sexual offspring, especially in late summer, when we observed that susceptible offspring resistotypes (RRSS⎵ and SSSS⎵, orange and yellow in Fig. 3C) were created from a parental population that consisted almost solely of resistant resistotypes (RRSR⎵ and RRRR⎵, blue in Fig. 3A). Remarkably, the most resistant resistotypes in the planktonic population were hardly seen in the offspring populations (RRRR⎵ in 2014 and 2015 (dark blue), RRRRR, RRRRS and RRSRR in 2017 (dark and bright blue)) (Fig. 3A and C).

Resting stages produced during the planktonic phase accumulate over the planktonic season, overwinter and hatch in the following spring. Pooling the resistotype data of the hatchlings from the sediment traps across the entire season and weighting resistotype frequencies by the abundance of resting stages in each sample therefore predicts the expected resistotype composition for the following spring cohort. These predictions match the resistotype composition of the planktonic population in spring very well for all three years (Fig. 4, Supplementary Figs. S3 and S4), indicating that our sediment traps are representative of these spring cohorts.

**Figure 4.**
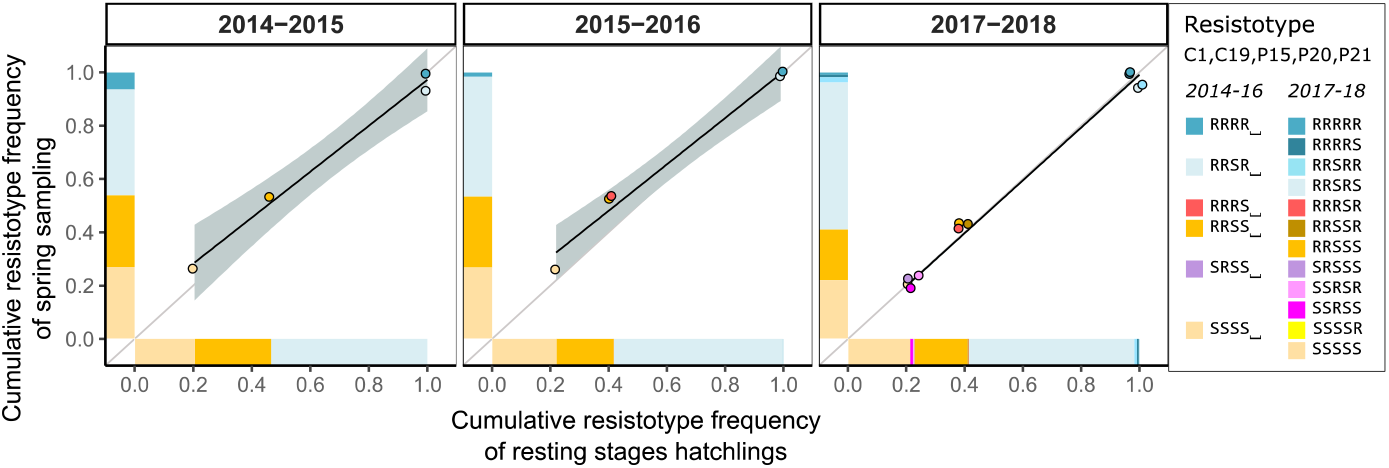
Scatter plot of resistotype frequencies of the hatchlings from the overwintering resting stages (collected in the sediment traps) against those of the *Daphnia magna* collected the following spring in the Aegelsee. The x-axis represents cumulated resistotype frequencies in the hatchlings from the sediment traps. These frequencies were calculated by weighing resistotype frequencies in the hatchling population by the relative number of resting stages produced at each sampling point. The y-axis represents cumulated resistotype frequencies in the first sample collected the following spring after the resting stages. Dots are plotted using jitter to reduce overlap. The grey line represents the *y* = *x* function and depicts an expected perfect match between both resistotype frequencies. The black line represents the fitted linear regression with 95% error as the grey area (not visible in the 2017-2018 panel because it is too small).

### Calculation of expected resistotype frequencies in resting stages

From previous work we know that dominance and epistasis are defining features of the inheritance of resistance to *P. ramosa* (Luijckx et al. 2013; Metzger et al. 2016; Bento et al. 2017; Bento et al. 2020; Ameline et al. 2021). To predict the role of sexual recombination in shaping resistance dynamics, we used an existing genetic model for resistance in our study population to calculate the expected resistotype frequencies in the offspring population at the time of resting stage production (sexual reproduction). These calculations require knowledge of allele frequencies at the resistance loci, which are unknown, but which we estimated using known resistotype distributions and assumptions. Although the published genetic model for resistance includes six loci (A to F), here we considered variation only at the B, C, D and E loci (Supplementary Fig. S5). The A and the F loci, known from other *D. magna* populations, seem to be monomorphic in the Aegelsee population. Variation at the B and the D loci seem strongly biased, with alleles B and d being rare: resistotypes determined by the “B-” and the “dd” genotypes, regardless of the genotype at other loci, only occur rarely, if at all, in our *D. magna* population (Fig. 1 and Ameline et al. 2021). Allele frequency at the C and E loci have been previously determined in a spring sample from the Aegelsee *D. magna* population (Ameline et al. 2021). Using these C- and E-loci allele frequencies within each resistotype and fixing the B and D loci to be “bbDD” genotype, we found that expected and observed resistotype frequencies match better than do other scenarios, e.g. equally distributed allele frequency at the C and E loci (Fig. 3C and E, Supplementary Figs. S6 and S7, Tables S1 and S2). Expected and observed resistotype frequency in the hatchling population match especially well in the first half of the season (Fig. 3C and E). In the second half of the season, however, we see a marked difference, with the presence of the abundant RRSS⎵ resistotype (about 25%) not predicted by the model (orange in Fig. 3C and E).

This discrepancy between predicted and observed resistotype frequencies in the second half of the season may have been due to a non-representative distribution of animals producing the sexual eggs (resting stages) at this time of the year. We tested this by collecting, in August 2020, *D. magna* samples and quantifying the resistotype distribution of females carrying resting stages, of males, and of a random sample of females. We found good correspondence between the random population samples, the sexual females, and the males (Supplementary Table S3), indicating that the animals reproducing sexually are a representative sample of the population with regards to resistotypes.

## Discussion

In this study we monitored the impact of seasonal epidemics of a bacterial pathogen (*Pasteuria ramosa*) on a natural zooplankton population (*Daphnia magna*), observing an increase in resistance phenotypes (resistotypes) every summer coinciding with epidemics, which has been shown to be driven by parasite mediated selection on two well-defined genomic regions (Ameline et al. 2021). Surprisingly, the frequencies of the different resistotypes among the spring hatchlings from the overwintering resting stages remained stable over the eight-year observation period, indicating an apparent absence of response to selection across years. In these cyclical dynamics there was an annual summer increase in mean resistance and an associated decline in the variance of resistance, followed by an over-winter decrease in resistance and increase in variance. We show that this cyclical pattern is shaped by both the seasonality of sexual reproduction and a genetic model for the inheritance of resistance involving dominance and epistasis.

### Repeated strong parasite-mediated selection in a natural population

The seasonal change in resistotype proportions in the *D. magna* Aegelsee population that we observed followed a consistent pattern across our eight-year study period: the ratio of resistant phenotypes increases during the planktonic phase of the host, coinciding with the *P. ramosa* epidemics (Fig. 1). Using material collected in the Aegelsee in 2014 and 2015, we previously confirmed that these resistotype patterns indeed resulted from parasite-mediated selection (Ameline et al. 2021), which has been shown to rapidly raise the frequency of resistant *Daphnia* (Little and Ebert 1999; Decaestecker et al. 2007; Duffy and Sivars-Becker 2007; Duncan and Little 2007) and other systems (Schmid-Hempel 2011, Morgan and Koskella 2017, Koskella 2018, Kurtz et al. 2016), although long-term monitoring of natural populations remains scarce (Laine 2009; González-Tortuero et al. 2016; Gibson et al. 2018).

We observed that this increase in resistance to different parasite genotypes occurred in the host population with some variation in time points, magnitude and speed (see seasonal increase in dark grey in Fig. 1C). Most notably, over five consecutive years, resistance to the sympatric *P. ramosa* isolate P20, which has been shown to play a major role in epidemics in our population (Ameline et al. 2021), increased from about 50% to 100% each year within two months around the peak of the epidemics. The results suggest that this is the case over the eight-year study period. Resistance to *P. ramosa* isolates C1 and C19 consistently increased throughout the planktonic phase each year, showing that resistance to these infectotypes may also be selected for in the host population. However, resistance was already high, about 75-80%, at the beginning of the planktonic phase, reaching about 100% by the end of the season. From 2016 to 2018, resistance to P15 and P21 increased as well, but decreased somewhat when parasite prevalence declined towards the end of the season, which was not the case for resistance to P20. This result might be explained by cost of resistance, as resistant genotypes lose their selective advantage once parasite prevalence declines below some level, making susceptible genotypes increase in frequency.

The *D. magna* census population size in the Aegelsee is estimated at over ten million individuals with an overwintering resting egg bank of about the same size. Together with the consistency of our observed pattern, these facts suggest that genetic drift plays only a very minor role in shaping the yearly resistotype dynamics in the Aegelsee *D. magna* population and that the observed changes are the result of adaptive evolution, which is not always the case in the *Daphnia* system. In a rock pool metapopulation of *D. magna*, for example, with populations of small effective and census sizes, gene flow and strong founder effects create more sensitivity to drift, with a relatively weak signal of adaptive evolution (Cabalzar et al. 2019).

Overall, we observed a highly repeatable, adaptive increase in resistance to all tested parasite infectotypes. Despite this increase in resistance, however, susceptibility to the parasite was created anew after a round of sexual reproduction in the spring cohort. Thus, we observed a stable long-term resistotype diversity across years (Fig. 1B).

### The effects of genetic recombination on resistotype composition

#### Decrease of population mean resistance

Because sexual reproduction is a prerequisite for resting stage formation in *D. magna*, we could decouple the effects of selection and genetic recombination on resistance in our host population. Overwintering happens only in the form of sexually produced resting stages, as planktonic individuals die off in early October due to the artificial warming of this sewage pond (Supplementary Fig. S1). We observed that the mean resistance of the spring hatchlings was much below the mean resistance from the previous fall (Fig 2B). While the strength of this effect was surprising to us, the observation that population regress back to the mean of the parental population before selection is well known (Falconer 1981; Lynch and Deng 1994; Otto 2009). Prolonged periods of asexual reproduction are believed to amplify this effect, which might explain the strength of genetic slippage observed in the studied population (Lynch and Deng 1994). However, despite regression back to the mean of the parental generation, it is usually expected that, under selection, the offspring mean will move away from the parents (response to selection). This did not happen in our population. We suggest that the combination of two aspects of this system may explain this finding: the timing of sexual reproduction and the genetic architecture of resistance.

We found that resting stages are not only produced at the end of the planktonic season, but already starting in somewhat irregular patterns during the season, with roughly two peaks, one early in the epidemic and one towards the end of the season (Fig. 3B). Resting stages produced at different times did not vary in fitness-related aspects (hatching rate, resting stages per ephippium, hatching time) (Supplementary Fig. S2), suggesting that their contribution to the next year spring cohort is approximately even. Thus, some of the sexual eggs (resting stages) that hatched in the spring were produced before selection acted on the parental generation, dampening the overall effect of selection on the spring cohort the following year. However, as typically more than 50% of the resting stages were produced after selection had increased resistance, this alone cannot explain the strong regression of mean resistance. We speculate that the genetic architecture underlying resistance may also contribute to this discrepancy.

#### Sex reestablishes resistance diversity

We phenotyped the hatchlings of the sexually produced resting stages collected in the sediment traps throughout the season. Early in the season, sexual offspring present approximately the same resistotype distribution as their planktonic parent population. Strikingly, however, resting stages collected late in the season show a markedly different resistotype from the planktonic host population at this time of the year. Namely, the parent population in the late season is composed of mainly resistant animals but produces about 50% susceptible offspring resistotypes (Fig. 3). Sexual recombination, coupled with a genetic architecture with epistasis and dominance, could create susceptible genotypes out of resistant ones.

To investigate how resistotype diversity is reestablished through sexual reproduction, or how resistant phenotypes can produce susceptible ones, we used a previously published genetic model for the inheritance of resistance in *D. magna* against *P. ramosa* infections (Metzger et al. 2016; Bento et al. 2020; Ameline et al. 2021) (described in Supplementary Fig. S5). This model allowed us to predict the resistotype frequencies of sexual offspring given a pool of parent resistotypes and their underlying genotypes. We then compared these predicted resistotype frequencies to those we observed among the resting stage hatchlings we collected throughout the season. In the early half of the season, our model worked rather well, with a slight discrepancy between the proportions of the RRSR⎵ resistotype (the model predicted more than we observed; light blue in Fig. 3C and E) and RRSS⎵ (the model predicted less than we observed, orange in Fig. 3C and E). We observed a stronger discrepancy between expected and observed resistotype distributions in the second half of the season. In particular, although P20-susceptible resistotypes (RRSS⎵, orange in Fig. 3C and E) are very common (about 25%) among the offspring resistotypes, according to our model, they should not be produced by a parent population where P20-resistant resistotypes dominate because resistance to P20 is recessive (Fig. 3, Supplementary Fig. S5). The genetic model of resistance displays strong epistasis and dominance, also influencing resistance to P20 (Bento et al. 2020; Ameline et al. 2021). Two loci, the C and the B loci, epistatically influence resistance to P20, but in the present case, this cannot explain the emergence of RRSS⎵ offspring from a parent population lacking RRSS⎵ individuals. Taken together, genetic recombination in this multi-locus system with epistasis and dominance seems likely to contribute to the maintenance of genetic diversity for resistance, although our genetic model is not complete and may be missing further epistatic interactions between the known loci or additional unknown loci.

Finally, the genetic model alone does not allow us to predict the frequencies of resistotypes after recombination without making assumptions about allele frequencies at these loci. We assumed allele frequencies derived from the overall observed resistotype diversity in the population and from previous estimates using genetic markers (see methods section and Ameline et al. 2021). We also assumed that allele frequencies underlying each resistotype did not change across the planktonic phase because we have no reason to expect changes in the frequencies of genotypes coding for the same resistotype. With more knowledge about the actual loci underlying the resistotypes, we may be able to predict resistotype frequencies better in the future. However, changing the assumptions for the allele frequencies to predict resistotype frequencies in sexual offspring did not produce enough of an effect to explain the mismatch between the fall parent generation and their sexual offspring.

### No evidence for pre-hatching or pre-zygotic selection related to resistotype

Pre-zygotic and/or pre-hatching selection could also possibly explain observed discrepancies between parent and offspring resistotypes. This could occur if different resistotypes in the planktonic population contributed unequally to sexual reproduction, producing males or resting stages differentially, or copulating at different rates. However, Orsini et al. (2016) suggest that the produced resting stages in *D. magna* population accurately represent the planktonic population, which agrees with an assessment in our study population indicating that the males and females that participate in sexual reproduction represent the resistotype distribution of the population well.

Another form of pre-zygotic selection could result from negative assortative mating that favors rare susceptible resistotypes. We cannot rule out an effect of assortative mating to explain the resistotype distribution in the offspring population. Assortative mating describes non-random mating between male and female genotypes or phenotypes. Positive assortative mating linked to body size and other traits has been found in a variety of animals, while negative assortative mating linked to immune genes (MHC) has been found in mice and humans (Wedekind et al. 1995; Chaix et al. 2008; Jiang et al. 2013). However, assortative mating in relation to immunity or resistance remains to be investigated in invertebrates, and as most population genetics models, including the present study, assume random mating, this is an important field for further study in the *D. magna* model species. Resistotype-dependent selection during diapause or hatching selection could also distort hatchling resistotype frequencies.

Finally, one may imagine that our ephippia sediment traps collected late in the season contained resting stages that had been produced earlier in the season and were re-suspended in the water column. However, several arguments speak against this. First, the pond does not contain fish, that may cause bioturbation. Second, the lake has no inflow, but only a very slow outflow, causing no detectable water movement. Third, at times when the *D. magna* population does not produce resting stages (the spring cohort in April), we find no resting stages in the sediment traps. Fourth, the same redistribution (in quantity and quality) would have needed to occur every year, as we observed the same patterns over three years. We thus conclude that water turbation is an unlikely explanation for the observed mismatch between resistotype distributions in the fall planktonic phase and the sexual stages it produced.

### The Red Queen theory for the maintenance of sex

Genetic recombination is a double-edged sword. On one side, it creates novel genotypes and phenotypes on which selection can act; on the other side, it may destroy coadapted gene complexes. At first sight, the latter seems to be the case on our study population because the recombinant offspring are less resistant than their parents. This seems to contradict the Red Queen hypothesis for the maintenance of sex (Jaenike 1978; Hamilton et al. 1990; Otto and Nuismer 2004; Salathé et al. 2008), which postulates that genetic recombination is advantageous for hosts because it overcomes the rapid adaptation of parasites to common host genotypes (Lively 2010; Gibson et al. 2018). Under this hypothesis, common hosts are expected to become over-infected, and rare hosts gain an advantage because at least some of them may be resistant to the common parasites.

In our study population, we did not have good information about the parasite infectotypes spreading throughout the year; however, we saw that prevalence was still high after mid-season, even though susceptible host resistotypes largely disappeared. This suggests that we may be missing infectotypes in our study population, at least in the late season. With better knowledge of the parasite population, we may witness yearly shifts in the frequency of parasite infectotypes in response to changing host resistotype frequency. In turn, host resistotypes may increase or decrease due to parasite selection. In this scenario, genetic recombination in the host in the second half of the season may not destroy coadapted gene complexes but rather destroy the host resistotypes that become the target of late-season parasites. While this coevolutionary scenario is consistent with the available data, it is a post-hoc explanation and requires further testing.

## Conclusion

In this study, we demonstrate parasite-mediated selection in a natural host population and elucidate the impact of sexual reproduction on resistance diversity. Our work stresses (i) the cyclical nature of host–parasite interactions, (ii) the very fast pace of parasite-driven changes in the host population and (iii) the fact that sexual recombination plays an important role in reshuffling allele combinations. Due to dominance and epistasis in the genetic architecture of resistance, this reshuffling resets the clock to the time before selection acted, rendering the response to selection zero. Although this is an extreme case of genetic slippage in response to sex, it is a powerful agent to maintain genetic diversity, which is a hallmark of resistance in natural populations of *Daphnia* and other animals, plants and bacteria (Altermatt and Ebert 2008; Desai and Currie 2015; Zhao et al. 2016; Cabalzar et al. 2019; Broniewski et al. 2020; Sallinen et al. 2020; White et al. 2020). As climatic seasonality seems to determine the dynamics of parasite resistance in our host population, and given the known impact of climate change on epidemics in the *D. magna*–*P. ramosa* system (Auld and Brand 2017), one may speculate that the dynamics in our study population may change in response to the predicted changes in climatic conditions and seasonality.

## Materials and methods

### *The* Daphnia magna – Pasteuria ramosa *system*

*Daphnia magna* Straus (Cladocera) is a freshwater planktonic crustacean that reproduces by cyclical parthenogenesis. Asexual females produce genetically identical (clones) diploid daughters or sons throughout the season. These females may switch to become sexual, and their haploid eggs need fertilization by males. Sexual eggs, which we call resting stages (precisely: embryos in developmental arrest) are produced, singly or in pairs, in a protective case (= ephippium) and require a resting period prior to hatching. All hatchlings from resting stages are asexually reproducing females. *Daphnia* filter-feed on planktonic algae and from the sediment surface, which is also how they ingest the transmission stages (= spores) of the bacterial parasite *Pasteuria ramosa* (Firmicutes: Bacillales). When infected by *P. ramosa, D. magna* take on a reddish coloration and increase in size (gigantism). Infection results in castration, reducing host reproductive success by 80% to 90%. Infected hosts die after six to ten weeks, releasing millions of long-lasting spores into the environment (Ebert et al. 1996; Ebert et al. 2016; Ben-Ami 2017).

### Temporal monitoring

Our study site was the Aegelsee pond near Frauenfeld, Switzerland, a fishless pond previously described in detail in Ameline et al. (2021), which contains a very large population of *D. magna*. To study the impact of the *P. ramosa* epidemics on the host, we sampled the *D. magna* population throughout its planktonic season (April to early October) for eight consecutive years, monitoring the frequencies of different resistance phenotypes (resistotypes) in the planktonic population. We also used traps to collect *D. magna* resting stages for three seasons and hatched them under semi-natural conditions the following spring. From 2011, a temperature logger was placed in the pond at a water depth of 0.5 m suspended from a buoy near the sampling spot. Water level was recorded at each sampling event.

#### Field work

Our first sample was collected in early October 2010. From 2011 to 2013, we collected approximately once a month, often a small sample size and without a standardized sampling protocol. From 2014 to 2018, we sampled the *D. magna* population using a standardized protocol every two to four weeks from early April to early October (more samples during the epidemic). Unless mentioned otherwise, all measurements were done at the deepest location close to the southern bank of the pond.

To monitor prevalence and the evolution of resistance, we sampled planktonic *D. magna* females at each collection date. We scooped the whole depth of the water column with a net (20-cm width and 1-mm mesh opening) to obtain several hundred animals. Samples were kept at 15 to 20 °C and transported to the laboratory and processed within four hours.

To sample the overwintering resting stages of the population, we collected surface sediment from five locations in the pond once in February 2014, before onset of the natural hatching season. This sample represents the overwintering resting population produced during the active season in 2013. To longitudinally sample the resting stages produced by the *D. magna* population across the season, we used five to nine sediment traps (vertically standing cylinders with 18-cm diameter and 0.4-mm mesh opening) placed on the lake bottom near the deepest part of the lake, and retrieved their content at each collection date during the planktonic season in 2014, 2015, 2017 and 2018. Collected *D. magna* resting stages were hatched in outside containers the following spring after overwintering at 4 °C in the dark. Each container contained a hundred ephippia per trap per timepoint and were monitored for several weeks. We collected hatchlings and cloned them in the laboratory. We measured the resistotype of 20 clonal lines (clones) per trap per timepoint, resulting in 100 clones per timepoint.

To obtain an estimate of *Daphnia* density, we used bottles to directly scoop the water from different depths three to five times (from 2011 to 2013). From 2014 we used a plankton net, performing ten vertical hauls from the bottom of the pond at the deepest point of the lake.

#### Analysis of field samples

The Aegelsee contains three *Daphnia* species: *D. magna*, *D. pulex* and *D. curvirostis*. The relative abundance of these species was measured in the laboratory by sorting and counting the density samples using a stereomicroscope. Because *D. pulex* and *D. curvirostis* have similar morphologies, we counted them together and inferred their relative proportions by determining the species in a random subset of 100 animals. We counted the number of males in a subset of 100 *D. magna*.

From each sample, we established clonal (iso-female) lines of about 100 *D. magna* to be used later for resistotype assessment. We estimated the prevalence of infection as described in (Ameline et al. 2021), and cured *P. ramosa* infections when they were observed, as otherwise cloning is not possible. *Pasteuria ramosa* is the only significant parasite in this population and was never observed to infect any species other than *D. magna* in this population.

We counted the *D. magna* ephippia retrieved from the traps and overwintered them at 4 °C in the dark. In the spring following the collection year (2014, 2015 and 2017), 20 to 100 ephippia (depending on how many were collected at a given sampling time point) from each sampling date were placed in 80-L containers filled with 30 L ADaM medium (Klüttgen et al. 1994, as modified in Ebert 1998). Containers were placed outdoors under direct sunlight and checked for hatchlings every second day. We recorded hatching dates and cloned hatchlings in the laboratory. We randomly chose 100 *D. magna* clones equally distributed among replicate traps at each sampling date to assess the resistotypes. To estimate hatching rate, we counted the number of resting stages per ephippium (zero, one or two) in a subset of ten to 20 ephippia that were not used for the hatching experiment, in at least two replicates for each collection date.

### Pasteuria ramosa *isolates*

In addition to the four *P. ramosa* isolates used previously (C1, C19, P15 and P20, Ameline et al. 2021), we isolated another strain of *P. ramosa,* P21,from our study population by exposing *D. magna* clones to suspended pond sediment. We took one infected female and serially passaged the bacteria from this female three times by infecting females of the same host clone. Spore production in the laboratory followed the protocol described by Luijckx et al. (2011).

### Resistotype assessment: the attachment test

We determined the resistance phenotype (resistotype) for each *D. magna* clone using the attachment test of Duneau et al. (2011). In short, early in the infection process, bacterial spores will attach to the foregut or the hindgut of susceptible host clones and penetrate the host’s body cavity. Spore attachment indicates host susceptibility (S), while absence of attachment indicates host resistance (R). We exposed each individual host to 8000 (C1, C19) or 10000 (P15, P20, P21) fluorescent spores and assessed attachment microscopically. Attachment was judged in each individual as yes or no. At least three replicates of each clone were used for each parasite isolate. Replicates are highly consistent in their attachment (Bento et al. 2017; Bento et al. 2020). For each host–parasite combination we obtained a consensus resistotype (R or S), based on the majority of the individual attachment tests. Across parasite strains, we defined the overall resistotype as the combination of resistance phenotypes to the five individual *P. ramosa* isolates in the following order: C1, C19, P15, P20 and P21 (e.g. a clone susceptible to all isolates will have the SSSSS resistotype). When resistance to a strain is not considered, we use the placeholder “⎵”, e.g. “RR⎵RR resistotype”. With time, we were able to include more parasite isolates: from 2010 to 2013, only the resistotypes to C1 and C19 were assessed. In 2014 and 2015, P15 and P20 were added, and all five *P. ramosa* isolates were tested from 2016.

To assess genetic slippage, we calculated the population mean resistance to *P. ramosa* for each sampling time. We assigned a resistance score to each resistotype ranging from zero to one to compare timepoints when we used different numbers of parasite isolates. For example, a host individual with a RRSRS resistotype was attributed a resistance score of 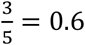.

### Hatching modelling

To predict resistotype frequencies of sexual offspring of the planktonic *D. magna* population-, we used the R-package “peas”, which generates predictions about the distribution of offspring genotypes and phenotypes in genetic crosses, based on specified systems of Mendelian inheritance (https://github.com/JanEngelstaedter/peas). We implemented the genetic model of resistance described in Ameline et al. (2021) in the *D. magna*–*P. ramosa* system for our study population. This model includes the genetic architecture of three loci (the B, C and E loci) that govern host resistance in our study population. The dominant allele at the B locus confers resistance (R) to C19 and susceptibility (S) to C1. The dominant allele at the C locus confers resistance to both the C1 and C19 *P. ramosa* strains, regardless of the genotype at the B locus (epistasis). The E locus contributes to resistance to P20. Resistance is dominant at the C locus (resistance to C1 and C19), whereas resistance is recessive at the E locus (resistance to P20). Homozygosity for the recessive allele at the B and C loci induces susceptibility to P20, regardless of the genotype at the E locus (epistasis). In the present study, we add the genetic architecture of the D locus to the model, which determines resistance to the P15 *P. ramosa* isolate (Bento et al. 2020). Implementation of the model is described in Supplementary Fig. S5 and Doc. S1. Implementing this model in the “peas” package, we calculated the expected resistance genotypes and phenotypes of sexual offspring of each possible mating among parent resistotypes. We assumed different allele frequency scenarios because the known resistotypes of the parents are not sufficient to estimate their exact genotype- and allele-frequencies, as some alleles can be hidden by dominance and epistasis. We then calculated the expected offspring resistotype frequencies over time corresponding to each of resting stage sample. If the genetic model accurately represents the biology of the system, the expected resistotype frequencies will match those found in the hatchlings from the sediment traps corresponding to the same sampling time. Detailed calculations are described in Supplementary Doc. S2 and Fig. S8.

### Statistical software

Unless otherwise stated, all statistical analyses and graphics were performed in the R software version 3.6.1 (R Core Team 2019). Graphics were edited in Inkscape v. 1.0.1 (https://inkscape.org/). Mean values are presented with standard error: mean ± se (Package RVAideMemoire v. 09-45-2, Hervé 2015). Packages used in R for package installation, data manipulation and graphics are the following: package development, documentation and installation: devtools v. 2.2.1 (Wickham, Hester, et al. 2019) and roxygen2 v. 6.1.1 (Wickham et al. 2018), data manipulation: dplyr v. 0.8.3 (Wickham, François, et al. 2019), tidyr v. 1.0.0 (Wickham and Henry 2019), tidyquant v. 0.5.8 (Dancho and Vaughan 2019), tidyverse v. 1.2.1 (Wickham 2017), xlsx v. 0.6.1 (Dragulescu and Arendt 2018), graphics: ggplot2 v. 3.3.0 (Wickham 2016), extrafont v. 0.17 (Chang 2014), scales v. 1.0.0 (Wickham 2018), cowplot v. 1.0.0 (Wilke 2019), gridExtra v. 2.3 (Auguie 2017), ggpubr v. 0.2.3 (Kassambara 2019), ggplotify v. 0.0.4 (Yu 2019), magick v. 2.2 (Ooms 2019), egg v. 0.4.5 (Auguie 2019), ggsci v. 2.9 (Xiao 2018) and png v. 0.1-7 (Urbanek 2013).

## Acknowledgments

We thank Yann Bourgeois, Samuel Pichon, Sabrina Gattis, Georgia Rouseti, Jürgen Hottinger, Urs Stiefel, Kristina Müller, Michelle Krebs and Dita Vizoso for help in the field and in the laboratory. Members of the Ebert group; Jonathon Stillman and Luís Teixeira provided valuable feedback on the study and the manuscript. This work was supported by the Swiss National Science Foundation (SNSF) (grant numbers 310030B_166677 and 310030_188887 to DE); the Freiwillige Akademische Gesellschaft (FAG) Basel to CA and the University of Basel to DE and CA.

## Additional information

### Author contributions

DE, JA and CA designed the study. JA conducted monitoring in 2010 to 2013, FV conducted monitoring and resting stage hatching in 2014, DE conducted monitoring in 2015 and CA conducted monitoring and resting stage hatching from 2016 to 2018. DE and ED monitored resistotype frequency and sexually reproducing animals in 2020 and 2021. JE developed the “peas” R package. CA analysed the data, designed figures and wrote the manuscript. All authors reviewed the manuscript.

