## Supplementary Material for "Genetic slippage after sex maintains diversity for parasite resistance in a natural host population"

#### Overview


| Section in main manuscript | Element | Description | Page |
|---|---|---|---|
| **Results** | | | |
| ***Seasonal epidemics*** | | | |
| | Fig. S1 | Environment and ecology | p. 2-3 |
| ***Selection and sexual reproduction*** | | | |
| | Fig. S2 | Hatching of planktonic resting stages | p. 4-5 |
| | Fig. S3 | Hatching of sediment-collected resting stages | p. 6 |
| | Fig. S4 | Overwintering resting stages | p. 7 |
| ***Calculation of expected resistotype frequencies in resting stages*** | | | |
| | Fig. S5 | BCDE genetic model | p. 8 |
| | | Description of allele frequency scenarios | p. 9 |
| | Fig. S6 | Results all scenarios | p. 10 |
| | Fig. S7 | Results best scenario | p. 11 |
| | Table S1 | Allele frequency scenarios | p. 12 |
| | Table S2 | Allele frequency best scenario | p. 13 |
| | Table S3 | Resistotype participation in sex | p. 14 |
| **Methods** | | | |
| ***Hatching modelling*** | | | |
| | Doc. S1 | "peas" genetic model | p. 15-17 |
| | Doc. S2 | Resistotype frequency calculations | p. 18-20 |
| | Fig. S8 | Summary of calculations | p. 21 |
| **References** | | | p. 22 |

### Results

##### *Seasonal epidemics - Environment and ecology in the Aegelsee*

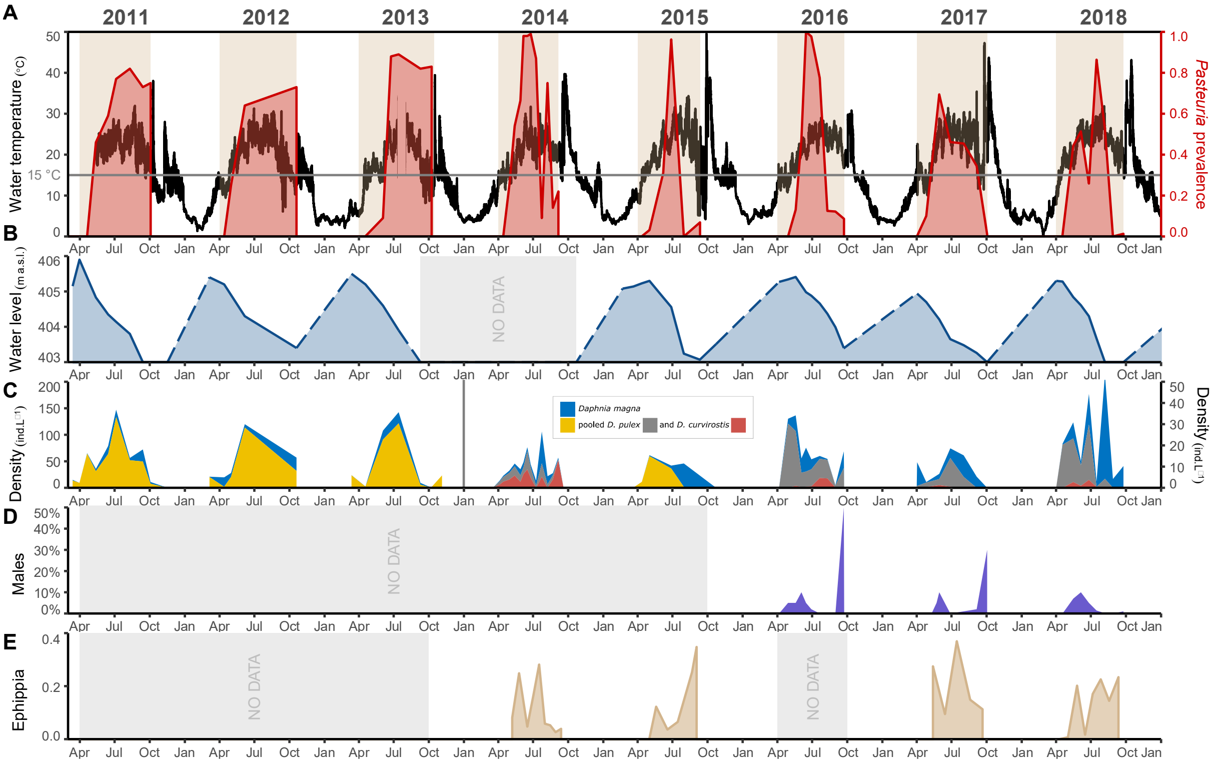


**Figure S1** Environmental conditions and sexual reproduction of *Daphnia magna* in the Aegelsee. **A:** Water temperature rise in the Aegelsee goes hand in hand with the appearance of the *Pasteuria ramosa* epidemics in the *D. magna* population. **Water temperature**: a temperature logger was installed in the pond from 2011 to 2018. No data is plotted in July 2013 because of vandalism of the data logger. The yearly temperature peaks in early October represent the release of warm ammoniacal condensation water in the pond. ***Pasteuria* prevalence**: red area plot represents *P. ramosa* prevalence in the *D. magna* population from 2011 to 2018. The grey horizontal line represents a water temperature threshold of 15 °C. When the water temperature rises above about 15 °C, the bacterial epidemics starts. **B:** Water level in the Aegelsee. Water level above sea level (a.s.l.) was read on a fixed floating device installed in the pond where the animals were sampled. We measured water level during the active season of the *D. magna*, from early April to early October. During this period, water level decreases progressively because of evaporation and agricultural and industrial use of the water. Solid line corresponds to data collected during the sampling period while dashed line corresponds to inferred data. Note that 403 m a.s.l. is the lower limit of the floating device. Every year in early October, warm condensation water is released in the pond. Over winter the pond is progressively filled up to a level of about 405 m a.s.l. **C:** *Daphnia* species density in the Aegelsee, measured as the number of individuals per liter of water. We sampled the water column during the active season, from early April to early October. From 2011 to 2013, pond water was directly sampled in 1-L bottles. From 2014 on, a plankton net was used. These two protocols created a four-fold magnitude difference between values obtained in 2011-2013 and 2014-2018, which we represent on distinct y-axes. *Daphnia* species were subsequently determined in the laboratory. As *D. pulex* and *D. curvirostis* are difficult to tell apart, a subset of 100 individuals of these two species was used to infer their respective densities in 2014, 2016, 2017 and 2018. They are pooled in the other years. In early October, warm ammoniacal condensation water is released in the pond, killing all plankton. No *Daphnia* overwinter in this population, neither do resting stages hatch before early April. **D:** *D. magna* male production in the Aegelsee. We counted males in a subset of 100 *D. magna* at each collection point from 2016 to 2018. **E:** *D. magna* ephippia production in the Aegelsee. Five to nine sediment traps were installed on the pond floor and retrieved at each collection date in 2014, 2015, 2017 and 2018. The y-axis represents ephippia number relative to the total number of ephippia counted in the season.

#### *Discussion about environmental and ecological variables*

We observe cyclical changes in different environmental variables. Every year, the *D. magna* population emerges when water temperature reaches about 12 °C (Fig. S1A). Temperature then increases to about 25 °C in summer, occasionally reaching peaks of 30 °C. In October, the warm ammoniacal condensation water is released in the pond, bringing temperature to 35-50 °C (Fig. S1A). The main increase in parasite prevalence occurs when water temperature rises above 15 °C (Fig. S1A). Water level in the Aegelsee is managed to make room for inflow of the condensation water in Fall. Therefore, every year the water level is lowered by about two meters over the course of the season. At its lowest level in late September, more than 80% of the pond sediments are exposed and the maximum water level is about one meter (Fig. S1B). *Daphnia* density shows irregular dynamics with a first peak typically in early summer, but further peaks may follow later. In most years, *D. magna* increases in relative frequency among all *Daphnia* species (Fig. S1C). We observe one or two peaks of *D. magna* male density during the season (Fig. S1D). We did not estimate the number of sexual females in the population, we instead collected resting stages in the sediment traps. Sexual egg counts cannot directly be compared with the frequencies of males, as they are time-shifted.

We observe a correlation between temperature cycles and *P. ramosa* epidemics in the *D. magna* Aegelsee population. Animals are observed to be infected by the bacteria as temperature rises above 15 °C every year in late April. We infer that epidemics are possibly influenced by water temperature, although the phenology of many other environmental factors may play a role. For example, longer day length, increased *Daphnia* density and lower water level (Fig. S1). It has been suggested that parasite-mediated selection in the *D. magna - P. ramosa* system is strongest at 20-25 °C (Mitchell et al. 2005; Vale et al. 2008). Given climate change model predictions of pond warming and longer seasons with temperatures above 15 °C, selection for resistance can thus be expected to intensify in our study population. Warming could also affect the evolution of stress tolerance, as exposure to the pathogen disrupts the host's ability to cope with thermal stress in this system (Hector et al. 2019). Environmental factors may also change the mode of selection: it has been shown that, under some temperature and food availability conditions, hosts in this system become more tolerant, thus potentially increasing parasite prevalence and slowing down coevolution (Vale et al. 2011). Parasite fitness may also be influenced by the interaction of genotype and environmental factors such as temperature and food availability (Vale and Little 2009, in a plant-parasite system: Laine 2007). Thus, while natural selection on resistance is precipitated on a high specificity of host-parasite interactions in the *D. magna* - *P. ramosa* system, it may also be linked to environmental conditions.

In the *D. magna* - *P. ramosa* system, host-parasite specificity is high, and spore attachment is not known to be influenced by environmental factors (Duneau et al. 2011; Luijckx et al. 2011). However, other host and parasite traits are influenced by the environment and there is intra-specific variability in how different genotypes respond to different environmental conditions (reviewed in Ebert et al. 2016). Temperature was found to influence infectivity and spore production in the parasite in the present system (Vale et al. 2008; Vale and Little 2009) and in the *D. dentifera* - *P. ramosa* system (Duffy and Hunsberger 2019). In the host, temperature was found to influence virulence in the present system (Mitchell et al. 2005) and filtering rate and parasite prevalence in a *D. laevis* - fungal host-pathogen system (Dallas and Drake 2016; Kirk et al. 2018; Kirk et al. 2019). Temperature also increased epidemic size in two mesocosm experiments, in the present system (Auld and Brand 2017) and in a *D. dentifera* - fungal parasite system (Shocket, Strauss, et al. 2018). This was explained in the latter *Daphnia* - fungus system by an increase of the transmission rate, composed of infectivity and foraging rate (Shocket, Vergara, et al. 2018). Nutrient availability increased tolerance of *D. magna* to *P. ramosa*, and increased spore production in the parasite, irrespective of temperature variations (Vale et al. 2011). Nutrient availability has also been shown to have a differential impact on fecundity and survival in distinct *D. magna* genotypes, leading to differential consequences of infection by a viral parasite (Reyserhove et al. 2017). Epidemiological variables such as prevalence, virulence, transmission rate and infection rate are thereby shaped by environmental variables (Hite and Cressler 2018).

***Selection and sexual reproduction***

**Figures S2 to S4**: The *Daphnia magna* overwintering resting stages in the Aegelsee

The *Daphnia magna* population in the Aegelsee goes through a cyclical pattern of resistotype (resistance phenotype) frequency. Resistant phenotypes increase in frequency over the course of the epidemics but resistotype diversity is created anew each spring via the hatching of the resting stages overwintering population. We collected and hatched ephippia laid in the water column by the planktonic population of *D. magna* throughout the active season in 2014, 2015, 2017 and 2018 using sediment traps (Figs. S2). We subsequently collected and hatched ephippia from surface sediment in winter 2014 as a representative sample of the spring *D. magna* cohort (Fig. S3). Figure S4 represents planktonic, ephippia and hatchling data together as a timeseries.

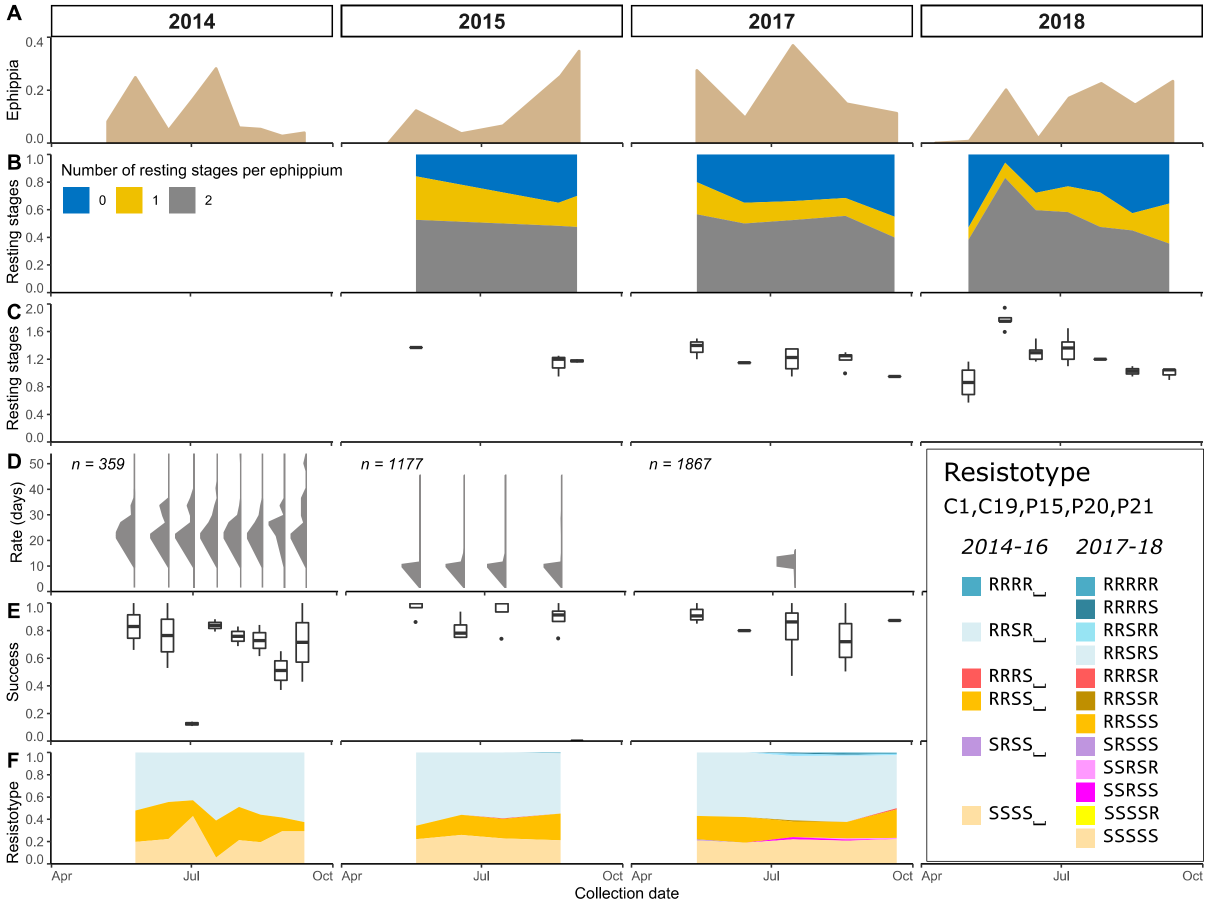


**Figure S2** Hatching of the *Daphnia magna* overwintering resting stages in the Aegelsee. Ephippia were collected in 2014, 2015, 2017 and 2018 during the active season. We used five to nine sediment traps at each collection date. Ephippia were subsequently stored at 4 °C to mimic the resting period and hatching was induced the following spring by placing the ephippia in outside containers. In 2014, the first ephippia sample was lost. In 2015, no ephippia hatched from the last sample because of exposure to the warm ammoniacal condensation water released into the pond at the end of the season. **A**: relative proportion of ephippia laid in the water column during the active season of *D. magna*. **B & C**: Number of resting stages per ephippium. On a subset of ephippia not used in the hatching experiment, we counted the number of resting stages present in the ephippial case. In ephippia collected in 2014, egg number was counted after the hatching experiment. This is why we do not plot it here, but we did use the values to infer hatching success (see E below). We checked ten to 20 ephippia for a minimum of two repeats at each collection date. **B**: Number of resting stages per ephippium, as the proportion of ephippia containing zero, one or two resting stages. **C**: Number of resting stages per ephippium, as the mean number of resting stages in the ephippial case. **D**: Hatching rate and pattern of collected ephippia. We induced hatching of the collected resting stages by putting 20 to 100 ephippia; depending on how many were collected; of each repeat of each collection date in outside containers. Resting stages collected in 2014, 2015 and 2017 were induced on 18 March 2015, 11 April 2016 and 29 March 2018, respectively. Hatching was monitored every second day. Vertical graphs represent the relative proportion of hatchlings over time after hatching induction. Each vertical graph represents ephippia from each collection date during the season. The different repeats for each collection date are pooled in one vertical graph. The y-axis (x-axis of the vertical graphs) is the number of days since hatching induction. In 2017, the total proportion of hatchlings across repeats and collection dates is presented. Resting stages collected on the last collection date of 2015 did not hatch because the warm ammoniacal condensation water had been released before our sampling. Variation across years might be due to the different hatching induction dates in the different years. **E**: Hatching success. We calculated hatching rate for each repeat for each collection date. We used the total number of hatchlings and the total number of resting stages induced, inferred from the subset of ephippia opened (B & C). For resting stages collected in 2014, only the number of remaining resting stages after hatching induction was counted. We inferred the total number of induced resting stages by adding the number of remaining resting stages to the total number of hatchlings. **F**: Resistotype (resistance phenotype) of hatched animals. Hatchlings were cloned in the laboratory to assess their resistotype. We tested 20 *D. magna* clonal lines (clones) for each repeat for each collection date, resulting in about 100 clones for each collection date.

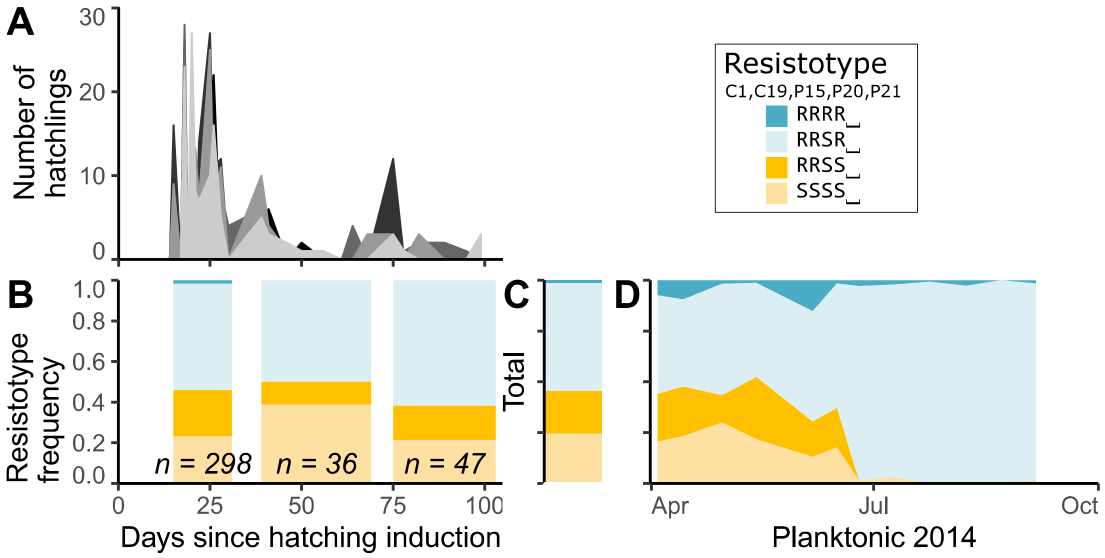


**Figure S3** *Daphnia magna* overwintering resting stages in the Aegelsee, collected in the sediment in winter 2014. The overwintering resting stages in winter 2014 were laid in the active season in 2013 and reflects the spring 2014 *D. magna* cohort. We collected five replicates of surface sediment in the pond in February 2014, before onset of the natural hatching season. A hundred ephippia from each replicate were placed in outdoor containers in late February 2014 and hatching was monitored every second day. **A**: Number of hatchlings over time after hatching induction. The five replicates are represented in different shades of grey. A total of 608 hatchlings were recorded. **B**: Resistotype (resistance phenotype) distribution of hatchlings over time after hatching induction. Hatchlings were put separately in jars to produce clonal lines. We measured resistotype on a subset of 381 randomly chosen clones. We represent resistotype frequency of hatched animals in three date intervals because of low sample sizes at some dates. The x-axis in A and B spans from day 0, the 20 February 2014 to day 103, the 3 June 2014. **C**: total resistotype proportions resulting from all 381 hatchlings. **D**: resistotype frequency of sampled *D. magna* in 2014.

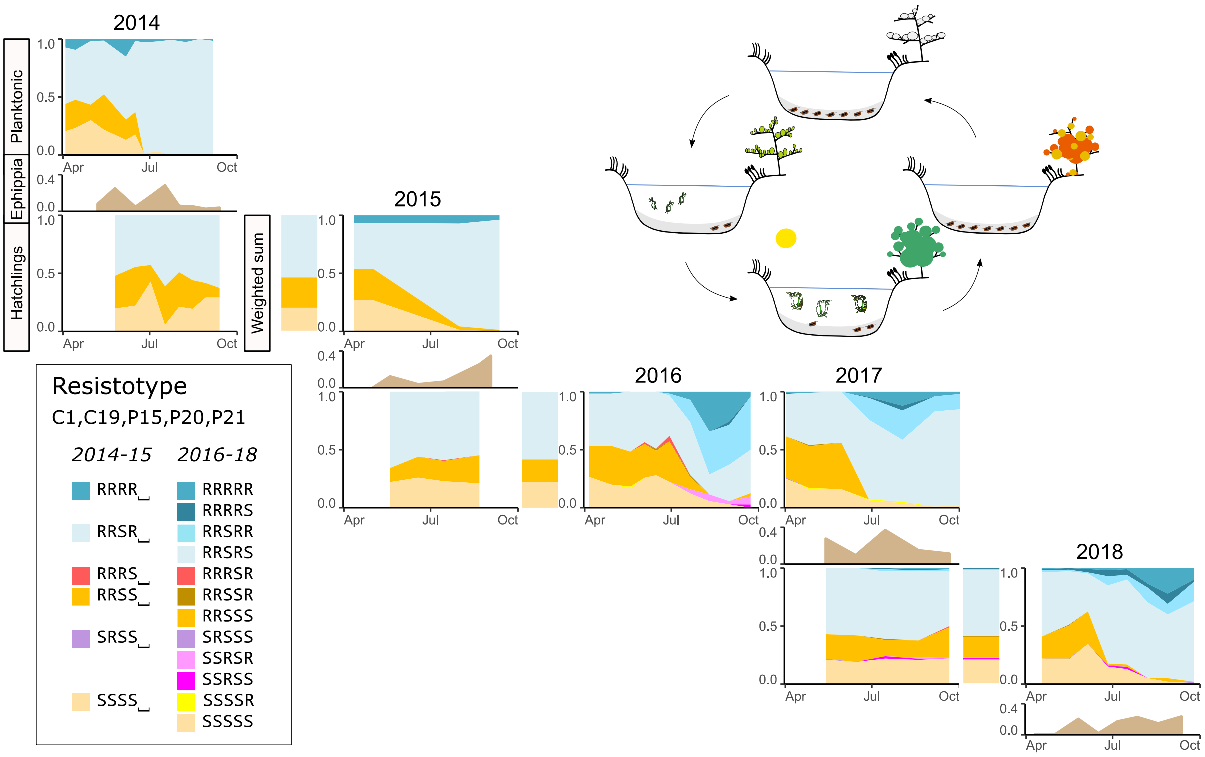


**Figure S4** *Daphnia magna* overwintering resting stages in the Aegelsee, collected in the water column throughout the active season. The spring *D. magna* cohort hatches from the resting stages present in the pond sediment. Throughout the active season (from early April to early October), *D. magna* reproduce asexually (clonal eggs) and sexually (fertilized resting stages). The resting stages create the overwintering population. In early October, warm condensation ammoniacal water is released in the pond, killing all plankton but not the resting stages. In winter, no ephippia hatch. **Planktonic**: Resistotype frequency in the *D. magna* population from 2014 to 2018. A large batch of animals was collected from early April to early October every 2-4 weeks to clone about 60 to 100 females. The resistotype is the full resistance phenotype to five *Pasteuria ramosa* isolates: C1, C19, P15, P20 and P21. Resistance and susceptibility are denoted as R and S, respectively. Note that in different years, different numbers of *P. ramosa* isolates were tested. We use the placeholder "␣" in the resistotype when a bacterial isolate was not tested. Resistance to P20 is highlighted because of its importance in the evolution of the host population (Ameline et al. 2021). **Ephippia**: relative number of *D. magna* ephippia laid in the pond. Five to nine ephippia traps were set up in 2014, 2015, 2017 and 2018 and collected every 2-4 weeks from early April to early October. We plot the mean number of ephippia per trap at each timepoint divided by the total mean number of ephippia laid during the whole year. Time on the x-axis represents the middle point between setup and collection of the traps. **Hatching**: *D. magna* resting stages collected in 2014, 2015 and 2017 were hatched in outside containers the following spring, after a resting period at 4 °C. In each container, 20 to 100 ephippia per trap per timepoint were placed, depending on how many were collected. Hatched animals were cloned in the laboratory. In *Daphnia*, hatchlings from resting stages are female, which allows to clone them. We measured the resistance phenotype (resistotype) of 20 clones per trap per timepoint, resulting in 100 clones per timepoint. **Weighted sum**: weighted sum of *D. magna* resistotype frequency from hatched ephippia. Total *D. magna* resistotype frequency from hatched ephippia weighted by the relative number of ephippia laid at each timepoint. The weighted sum of resistotype frequency represents the overwintering resting stages. In 2014, the first ephippia sample was lost. In 2015, no ephippia hatched from the last sample because of exposure to the warm ammoniacal condensation water released into the pond at the end of the season.

##### *Calculation of expected resistotype frequencies in resting stages*

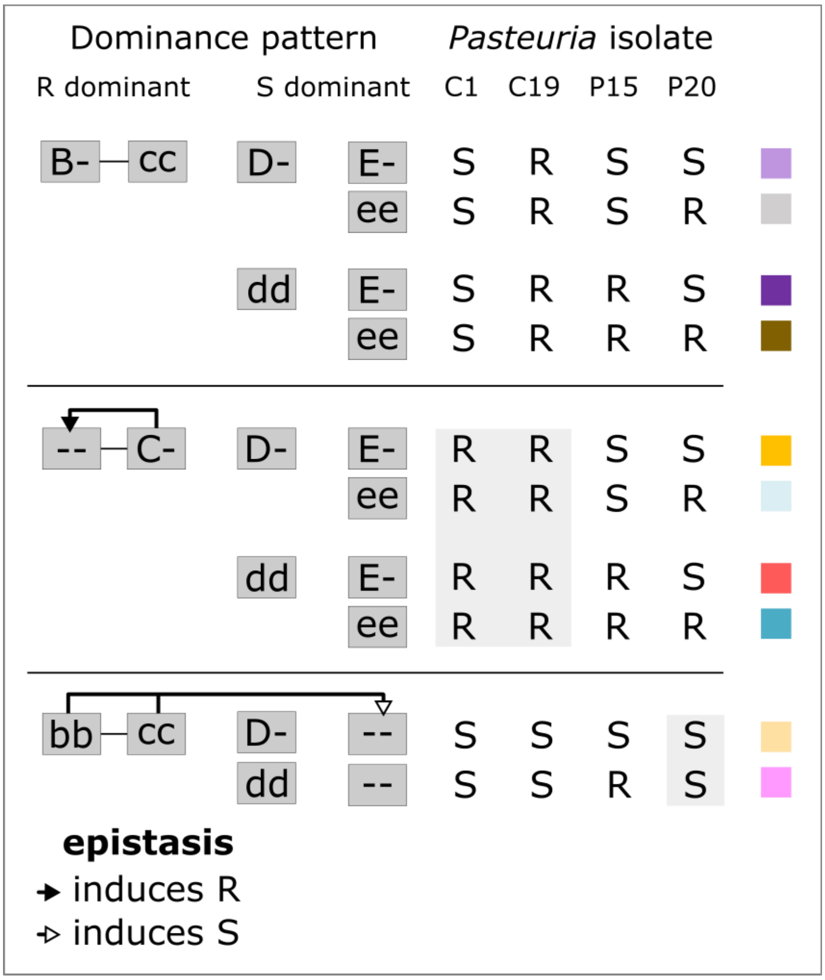


**Figure S5** genetic model of resistance in the Aegelsee. The model includes resistotypes to C1, C19, P15 and P20 *Pasteuria ramosa* isolates. Resistance to C1 and C19 determined by the ABC-cluster was described in Metzger et al. (2016). The dominant allele at the B-locus induces resistance (R) to C19 and susceptibility (S) to C1. The dominant allele at the C-locus confers resistance to both C1 and C19 *P. ramosa* isolates, regardless of the genotype at the B-locus. Variation at the A-locus is not considered here as the recessive allele at this locus is believed to be fixed in the population (Ameline et al. 2021). The D-locus determines resistance to P15 (Bento et al. 2020). The E-locus determines resistance to P20 (Ameline et al. 2021). Resistance is dominant at the B- and C-loci (resistance to C1 and C19) whereas resistance is recessive at the D- and E-loci (resistance to P15 and P20, respectively). Recessive homozygosity at the B- and C-locus induces susceptibility to P20, regardless of the genotype at the E-locus (Ameline et al. 2021). Hence the epistasis can only be observed phenotypically in "bbccee" (SS␣S) offspring. If the epistatic relationship is not present in this case, the observed phenotype would be SS␣R. Such SS␣R individuals were never observed in the population. Resistotypes determined by the "B-" and the "dd" genotype, regardless of the genotype at other loci, are very rare or do not occur in the *D. magna* population. We assume the B- and the d-alleles are rare in the population and might induce poor fitness. We implement this genetic model of resistance in the *D. magna* - *P. ramosa* system in the "peas" R package (Doc. S1). We subsequently test the model using resting stages hatching data (Figs. S6 and S7, Tables S1 and S2).

**Figs S6 and S7, Tables S1 and S2**: Predicted resistotype frequency resulting from resting stages hatching in the Aegelsee

Using the genetic model of resistance inheritance in the *Daphnia magna* - *Pasteuria ramosa* system, we calculate predicted resistotype frequency resulting from hatching of *D. magna* resting stages, or ephippia, produced throughout the active season. We compare this expected resistotype frequency to the observed resistotype frequency obtained from hatching of field-collected resting stages. The genetic model and calculations are described in Fig. S5, Doc. S1, Doc. S2 and Fig. S8. In short, we use three input datasets: (i) the longitudinally observed resistotype frequency, from the F0 generation performing sexual reproduction during the active season, (ii) the predicted F1 resistotype segregation given by the genetic model and (iii) the genotype distribution within resistotypes in the *D. magna* population, the F0 generation, described here. Because we use phenotype distribution data, we input genotype distribution within each phenotype. **Figs. S6 and S7**: observed vs. expected resistotype frequency resulting from resting stages hatching. We calculate expected resistotype frequency according to different allele frequency scenarios in the *D. magna* population. **Tables S1 and S2**: Allele frequency scenarios in the *D. magna* population. We use the list of possible genotypes and their corresponding resistotypes given by the genetic model of resistance in the system. In each scenario, we fix an allele at one or several loci and we equally distribute genotype proportions among the other loci. In resistotypes where it is not possible to fix the allele, we equally distribute genotype proportions among heterozygous genotypes or among homozygous genotypes for the alternative allele when this is the only possible genotype determining the resistotype. **Table S2** presents the “bbDD” scenario, additionally implemented with observed allele frequency at the C- and E-loci. Observed C- and E-loci allele frequency were measured in spring sample (Ameline et al. 2021).

- **scenario “bbDD”**: the “bb” and “DD” genotypes are fixed in all possible resistotypes.
- **scenario “bb”**: the “bb” genotype is fixed in all possible resistotypes.
- **scenario “BB”**: the “BB” genotype is fixed in all possible resistotypes.
- **scenario “CC”**: the “CC” genotype is fixed in all possible resistotypes.
- **scenario “DD”**: the “DD” genotype is fixed in all possible resistotypes.
- **scenario “ee”**: the “ee” genotype is fixed in all possible resistotypes.
- **scenario “EE”**: the “EE” genotype is fixed in all possible resistotypes.
- **scenario “hetero”**: all loci show heterozygous genotype in all possible resistotypes.

*Note*: we use four-letter resistotype because the genetic model of resistance includes resistance to the four *P. ramosa* isolates C1, C19, P15 and P20.

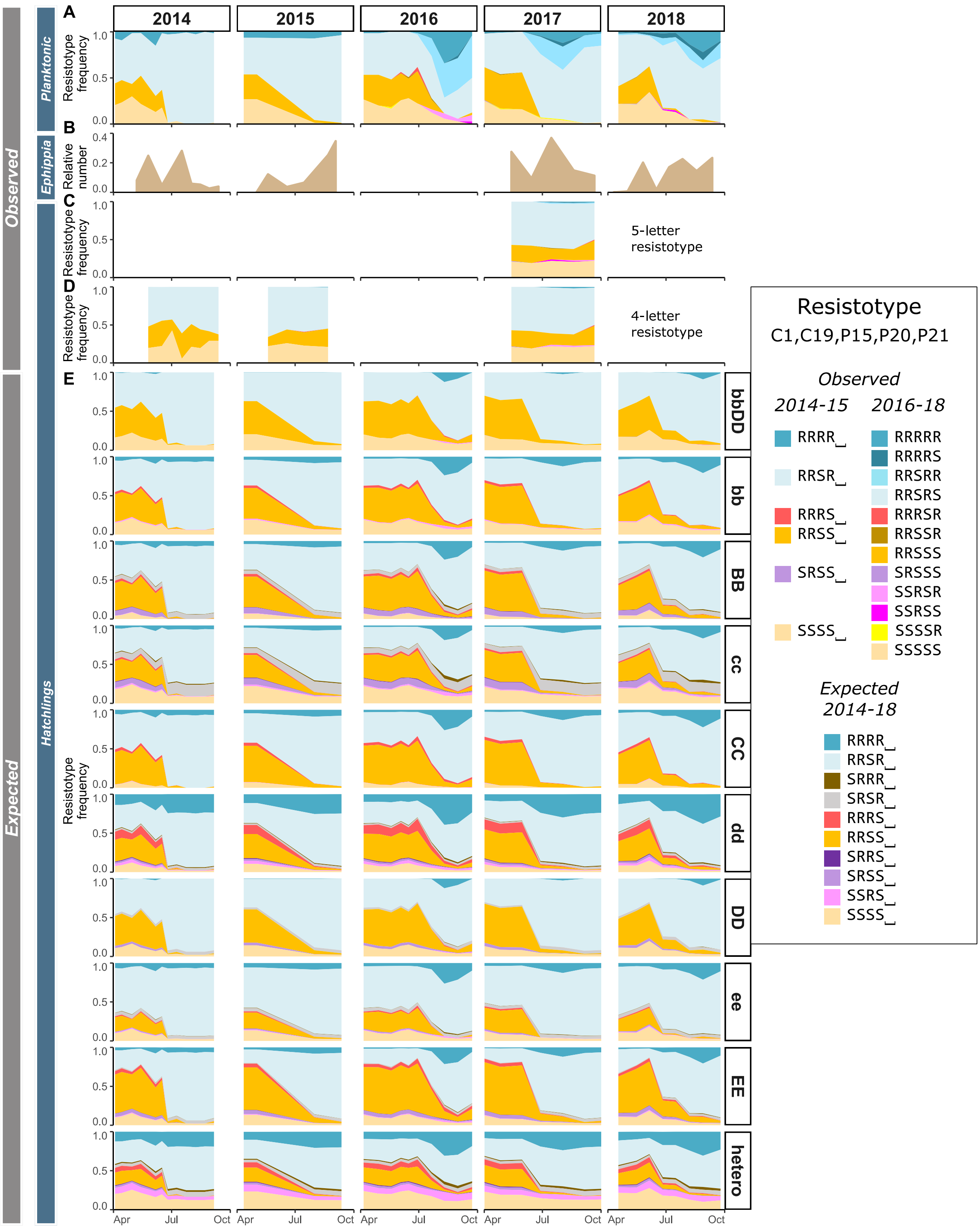


**Figure S5** Expected resistotype frequency resulting from resting stages, or ephippia, laid throughout the active season of *Daphnia magna*. We use the genetic model of resistance inheritance in the *Daphnia magna* - *Pasteuria ramosa* system, presented in Fig. S5. Allele frequency scenarios are detailed in Table S1.

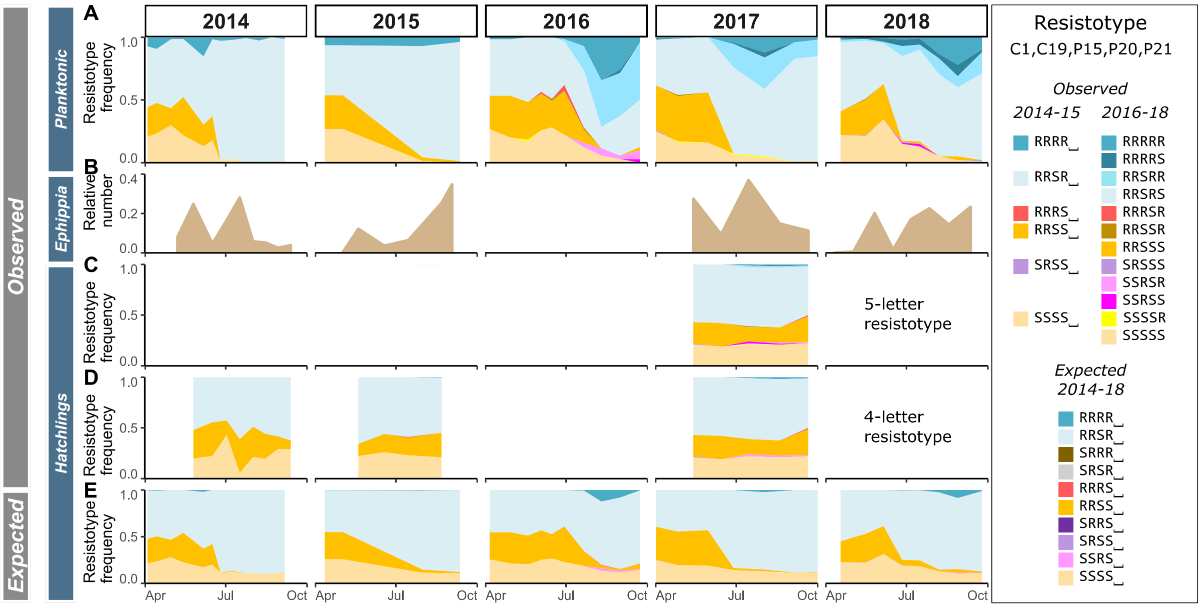


**Figure S6** Expected resistotype frequency resulting from resting stages produced throughout the active season of *Daphnia magna*. We use the genetic model of resistance inheritance in the *Daphnia magna - Pasteuria ramosa* system, presented in Fig. S5. This allele frequency scenario produces an expected resistotype distribution that fits best the observed one. The scenario is detailed in Table S2.

**Table S1** Genotype distribution scenarios in the Aegelsee *Daphnia magna* population. In each scenario we fix an allele at one or two loci and we equally distribute proportions in the remaining possible genotypes, within each resistotype. In resistotypes where it is not possible to fix the allele, we equally distribute genotype proportions among heterozygous genotypes or among homozygous genotypes for the alternative allele when this is the only possible genotype determining the resistotype. The genetic model described in Fig. S5 provided the list of possible genotypes and their corresponding resistotypes. The different scenarios are described above. This table corresponds to the “freq” table described in Supplementary Doc. S2 and Fig. S8.

<table>
<tr><th rowspan="2">pheno</th><th rowspan="2">geno</th><th colspan="10">Scenarios of genotype distribution within resistotypes</th></tr>
<tr><th>bbDD</th><th>bb</th><th>BB</th><th>cc</th><th>CC</th><th>dd</th><th>DD</th><th>ee</th><th>EE</th><th>hetero</th></tr>
<tr><td>RRRR</td><td>bb~Cc | dd | ee</td><td>1/2</td><td>1/2</td><td>0</td><td>1/3</td><td>0</td><td>1/6</td><td>1/6</td><td>1/6</td><td>1/6</td><td>0</td></tr>
<tr><td>RRRR</td><td>Bb~Cc | dd | ee</td><td>0</td><td>0</td><td>0</td><td>1/3</td><td>0</td><td>1/6</td><td>1/6</td><td>1/6</td><td>1/6</td><td>1</td></tr>
<tr><td>RRRR</td><td>BB~Cc | dd | ee</td><td>0</td><td>0</td><td>1/2</td><td>1/3</td><td>0</td><td>1/6</td><td>1/6</td><td>1/6</td><td>1/6</td><td>0</td></tr>
<tr><td>RRRR</td><td>bb~CC | dd | ee</td><td>1/2</td><td>1/2</td><td>0</td><td>0</td><td>1/3</td><td>1/6</td><td>1/6</td><td>1/6</td><td>1/6</td><td>0</td></tr>
<tr><td>RRRR</td><td>Bb~CC | dd | ee</td><td>0</td><td>0</td><td>0</td><td>0</td><td>1/3</td><td>1/6</td><td>1/6</td><td>1/6</td><td>1/6</td><td>0</td></tr>
<tr><td>RRRR</td><td>BB~CC | dd | ee</td><td>0</td><td>0</td><td>1/2</td><td>0</td><td>1/3</td><td>1/6</td><td>1/6</td><td>1/6</td><td>1/6</td><td>0</td></tr>
<tr><td>RRSR</td><td>bb~Cc | Dd | ee</td><td>0</td><td>1/4</td><td>0</td><td>1/6</td><td>0</td><td>1/6</td><td>0</td><td>1/12</td><td>1/12</td><td>0</td></tr>
<tr><td>RRSR</td><td>Bb~Cc | Dd | ee</td><td>0</td><td>0</td><td>0</td><td>1/6</td><td>0</td><td>1/6</td><td>0</td><td>1/12</td><td>1/12</td><td>1</td></tr>
<tr><td>RRSR</td><td>BB~Cc | Dd | ee</td><td>0</td><td>0</td><td>1/4</td><td>1/6</td><td>0</td><td>1/6</td><td>0</td><td>1/12</td><td>1/12</td><td>0</td></tr>
<tr><td>RRSR</td><td>bb~CC | Dd | ee</td><td>0</td><td>1/4</td><td>0</td><td>0</td><td>1/6</td><td>1/6</td><td>0</td><td>1/12</td><td>1/12</td><td>0</td></tr>
<tr><td>RRSR</td><td>Bb~CC | Dd | ee</td><td>0</td><td>0</td><td>0</td><td>0</td><td>1/6</td><td>1/6</td><td>0</td><td>1/12</td><td>1/12</td><td>0</td></tr>
<tr><td>RRSR</td><td>BB~CC | Dd | ee</td><td>0</td><td>0</td><td>1/4</td><td>0</td><td>1/6</td><td>1/6</td><td>0</td><td>1/12</td><td>1/12</td><td>0</td></tr>
<tr><td>RRSR</td><td>bb~Cc | DD | ee</td><td>1/2</td><td>1/4</td><td>0</td><td>1/6</td><td>0</td><td>0</td><td>1/6</td><td>1/12</td><td>1/12</td><td>0</td></tr>
<tr><td>RRSR</td><td>Bb~Cc | DD | ee</td><td>0</td><td>0</td><td>0</td><td>1/6</td><td>0</td><td>0</td><td>1/6</td><td>1/12</td><td>1/12</td><td>0</td></tr>
<tr><td>RRSR</td><td>BB~Cc | DD | ee</td><td>0</td><td>0</td><td>1/4</td><td>1/6</td><td>0</td><td>0</td><td>1/6</td><td>1/12</td><td>1/12</td><td>0</td></tr>
<tr><td>RRSR</td><td>bb~CC | DD | ee</td><td>1/2</td><td>1/4</td><td>0</td><td>0</td><td>1/6</td><td>0</td><td>1/6</td><td>1/12</td><td>1/12</td><td>0</td></tr>
<tr><td>RRSR</td><td>Bb~CC | DD | ee</td><td>0</td><td>0</td><td>0</td><td>0</td><td>1/6</td><td>0</td><td>1/6</td><td>1/12</td><td>1/12</td><td>0</td></tr>
<tr><td>RRSR</td><td>BB~CC | DD | ee</td><td>0</td><td>0</td><td>1/4</td><td>0</td><td>1/6</td><td>0</td><td>1/6</td><td>1/12</td><td>1/12</td><td>0</td></tr>
<tr><td>SRRR</td><td>Bb~cc | dd | ee</td><td>1</td><td>1</td><td>0</td><td>1/2</td><td>1/2</td><td>1/2</td><td>1/2</td><td>1/2</td><td>1/2</td><td>1</td></tr>
<tr><td>SRRR</td><td>BB~cc | dd | ee</td><td>0</td><td>0</td><td>1</td><td>1/2</td><td>1/2</td><td>1/2</td><td>1/2</td><td>1/2</td><td>1/2</td><td>0</td></tr>
<tr><td>SRSR</td><td>Bb~cc | Dd | ee</td><td>0</td><td>1/2</td><td>0</td><td>1/4</td><td>1/4</td><td>1/2</td><td>0</td><td>1/4</td><td>1/4</td><td>1</td></tr>
<tr><td>SRSR</td><td>BB~cc | Dd | ee</td><td>0</td><td>0</td><td>1/2</td><td>1/4</td><td>1/4</td><td>1/2</td><td>0</td><td>1/4</td><td>1/4</td><td>0</td></tr>
<tr><td>SRSR</td><td>Bb~cc | DD | ee</td><td>1</td><td>1/2</td><td>0</td><td>1/4</td><td>1/4</td><td>0</td><td>1/2</td><td>1/4</td><td>1/4</td><td>0</td></tr>
<tr><td>SRSR</td><td>BB~cc | DD | ee</td><td>0</td><td>0</td><td>1/2</td><td>1/4</td><td>1/4</td><td>0</td><td>1/2</td><td>1/4</td><td>1/4</td><td>0</td></tr>
<tr><td>RRRS</td><td>bb~Cc | dd | Ee</td><td>1/4</td><td>1/4</td><td>0</td><td>1/6</td><td>0</td><td>1/12</td><td>1/12</td><td>1/6</td><td>0</td><td>0</td></tr>
<tr><td>RRRS</td><td>Bb~Cc | dd | Ee</td><td>0</td><td>0</td><td>0</td><td>1/6</td><td>0</td><td>1/12</td><td>1/12</td><td>1/6</td><td>0</td><td>1</td></tr>
<tr><td>RRRS</td><td>BB~Cc | dd | Ee</td><td>0</td><td>0</td><td>1/4</td><td>1/6</td><td>0</td><td>1/12</td><td>1/12</td><td>1/6</td><td>0</td><td>0</td></tr>
<tr><td>RRRS</td><td>bb~CC | dd | Ee</td><td>1/4</td><td>1/4</td><td>0</td><td>0</td><td>1/6</td><td>1/12</td><td>1/12</td><td>1/6</td><td>0</td><td>0</td></tr>
<tr><td>RRRS</td><td>Bb~CC | dd | Ee</td><td>0</td><td>0</td><td>0</td><td>0</td><td>1/6</td><td>1/12</td><td>1/12</td><td>1/6</td><td>0</td><td>0</td></tr>
<tr><td>RRRS</td><td>BB~CC | dd | Ee</td><td>0</td><td>0</td><td>1/4</td><td>0</td><td>1/6</td><td>1/12</td><td>1/12</td><td>1/6</td><td>0</td><td>0</td></tr>
<tr><td>RRRS</td><td>bb~Cc | dd | EE</td><td>1/4</td><td>1/4</td><td>0</td><td>1/6</td><td>0</td><td>1/12</td><td>1/12</td><td>0</td><td>1/6</td><td>0</td></tr>
<tr><td>RRRS</td><td>Bb~Cc | dd | EE</td><td>0</td><td>0</td><td>0</td><td>1/6</td><td>0</td><td>1/12</td><td>1/12</td><td>0</td><td>1/6</td><td>0</td></tr>
<tr><td>RRRS</td><td>BB~Cc | dd | EE</td><td>0</td><td>0</td><td>1/4</td><td>1/6</td><td>0</td><td>1/12</td><td>1/12</td><td>0</td><td>1/6</td><td>0</td></tr>
<tr><td>RRRS</td><td>bb~CC | dd | EE</td><td>1/4</td><td>1/4</td><td>0</td><td>0</td><td>1/6</td><td>1/12</td><td>1/12</td><td>0</td><td>1/6</td><td>0</td></tr>
<tr><td>RRRS</td><td>Bb~CC | dd | EE</td><td>0</td><td>0</td><td>0</td><td>0</td><td>1/6</td><td>1/12</td><td>1/12</td><td>0</td><td>1/6</td><td>0</td></tr>
<tr><td>RRRS</td><td>BB~CC | dd | EE</td><td>0</td><td>0</td><td>1/4</td><td>0</td><td>1/6</td><td>1/12</td><td>1/12</td><td>0</td><td>1/6</td><td>0</td></tr>
<tr><td>RRSS</td><td>bb~Cc | Dd | Ee</td><td>0</td><td>1/8</td><td>0</td><td>1/12</td><td>0</td><td>1/12</td><td>0</td><td>1/12</td><td>0</td><td>0</td></tr>
<tr><td>RRSS</td><td>Bb~Cc | Dd | Ee</td><td>0</td><td>0</td><td>0</td><td>1/12</td><td>0</td><td>1/12</td><td>0</td><td>1/12</td><td>0</td><td>1</td></tr>
<tr><td>RRSS</td><td>BB~Cc | Dd | Ee</td><td>0</td><td>0</td><td>1/8</td><td>1/12</td><td>0</td><td>1/12</td><td>0</td><td>1/12</td><td>0</td><td>0</td></tr>
<tr><td>RRSS</td><td>bb~CC | Dd | Ee</td><td>0</td><td>1/8</td><td>0</td><td>0</td><td>1/12</td><td>1/12</td><td>0</td><td>1/12</td><td>0</td><td>0</td></tr>
<tr><td>RRSS</td><td>Bb~CC | Dd | Ee</td><td>0</td><td>0</td><td>0</td><td>0</td><td>1/12</td><td>1/12</td><td>0</td><td>1/12</td><td>0</td><td>0</td></tr>
<tr><td>RRSS</td><td>BB~CC | Dd | Ee</td><td>0</td><td>0</td><td>1/8</td><td>0</td><td>1/12</td><td>1/12</td><td>0</td><td>1/12</td><td>0</td><td>0</td></tr>
<tr><td>RRSS</td><td>bb~Cc | DD | Ee</td><td>1/4</td><td>1/8</td><td>0</td><td>1/12</td><td>0</td><td>0</td><td>1/12</td><td>1/12</td><td>0</td><td>0</td></tr>
<tr><td>RRSS</td><td>Bb~Cc | DD | Ee</td><td>0</td><td>0</td><td>0</td><td>1/12</td><td>0</td><td>0</td><td>1/12</td><td>1/12</td><td>0</td><td>0</td></tr>
<tr><td>RRSS</td><td>BB~Cc | DD | Ee</td><td>0</td><td>0</td><td>1/8</td><td>1/12</td><td>0</td><td>0</td><td>1/12</td><td>1/12</td><td>0</td><td>0</td></tr>
<tr><td>RRSS</td><td>bb~CC | DD | Ee</td><td>1/4</td><td>1/8</td><td>0</td><td>0</td><td>1/12</td><td>0</td><td>1/12</td><td>1/12</td><td>0</td><td>0</td></tr>
<tr><td>RRSS</td><td>Bb~CC | DD | Ee</td><td>0</td><td>0</td><td>0</td><td>0</td><td>1/12</td><td>0</td><td>1/12</td><td>1/12</td><td>0</td><td>0</td></tr>
<tr><td>RRSS</td><td>BB~CC | DD | Ee</td><td>0</td><td>0</td><td>1/8</td><td>0</td><td>1/12</td><td>0</td><td>1/12</td><td>1/12</td><td>0</td><td>0</td></tr>
<tr><td>RRSS</td><td>bb~Cc | Dd | EE</td><td>0</td><td>1/8</td><td>0</td><td>1/12</td><td>0</td><td>1/12</td><td>0</td><td>0</td><td>1/12</td><td>0</td></tr>
<tr><td>RRSS</td><td>Bb~Cc | Dd | EE</td><td>0</td><td>0</td><td>0</td><td>1/12</td><td>0</td><td>1/12</td><td>0</td><td>0</td><td>1/12</td><td>0</td></tr>
<tr><td>RRSS</td><td>BB~Cc | Dd | EE</td><td>0</td><td>0</td><td>1/8</td><td>1/12</td><td>0</td><td>1/12</td><td>0</td><td>0</td><td>1/12</td><td>0</td></tr>
<tr><td>RRSS</td><td>bb~CC | Dd | EE</td><td>0</td><td>1/8</td><td>0</td><td>0</td><td>1/12</td><td>1/12</td><td>0</td><td>0</td><td>1/12</td><td>0</td></tr>
<tr><td>RRSS</td><td>Bb~CC | Dd | EE</td><td>0</td><td>0</td><td>0</td><td>0</td><td>1/12</td><td>1/12</td><td>0</td><td>0</td><td>1/12</td><td>0</td></tr>
<tr><td>RRSS</td><td>BB~CC | Dd | EE</td><td>0</td><td>0</td><td>1/8</td><td>0</td><td>1/12</td><td>1/12</td><td>0</td><td>0</td><td>1/12</td><td>0</td></tr>
<tr><td>RRSS</td><td>bb~Cc | DD | EE</td><td>1/4</td><td>1/8</td><td>0</td><td>1/12</td><td>0</td><td>0</td><td>1/12</td><td>0</td><td>1/12</td><td>0</td></tr>
<tr><td>RRSS</td><td>Bb~Cc | DD | EE</td><td>0</td><td>0</td><td>0</td><td>1/12</td><td>0</td><td>0</td><td>1/12</td><td>0</td><td>1/12</td><td>0</td></tr>
<tr><td>RRSS</td><td>BB~Cc | DD | EE</td><td>0</td><td>0</td><td>1/8</td><td>1/12</td><td>0</td><td>0</td><td>1/12</td><td>0</td><td>1/12</td><td>0</td></tr>
<tr><td>RRSS</td><td>bb~CC | DD | EE</td><td>1/4</td><td>1/8</td><td>0</td><td>0</td><td>1/12</td><td>0</td><td>1/12</td><td>0</td><td>1/12</td><td>0</td></tr>
<tr><td>RRSS</td><td>Bb~CC | DD | EE</td><td>0</td><td>0</td><td>0</td><td>0</td><td>1/12</td><td>0</td><td>1/12</td><td>0</td><td>1/12</td><td>0</td></tr>
<tr><td>RRSS</td><td>BB~CC | DD | EE</td><td>0</td><td>0</td><td>1/8</td><td>0</td><td>1/12</td><td>0</td><td>1/12</td><td>0</td><td>1/12</td><td>0</td></tr>
<tr><td>SRRS</td><td>Bb~cc | dd | Ee</td><td>1/2</td><td>1/2</td><td>0</td><td>1/4</td><td>1/4</td><td>1/4</td><td>1/4</td><td>1/2</td><td>0</td><td>1</td></tr>
<tr><td>SRRS</td><td>BB~cc | dd | Ee</td><td>0</td><td>0</td><td>1/2</td><td>1/4</td><td>1/4</td><td>1/4</td><td>1/4</td><td>1/2</td><td>0</td><td>0</td></tr>
<tr><td>SRRS</td><td>Bb~cc | dd | EE</td><td>1/2</td><td>1/2</td><td>0</td><td>1/4</td><td>1/4</td><td>1/4</td><td>1/4</td><td>0</td><td>1/2</td><td>0</td></tr>
<tr><td>SRRS</td><td>BB~cc | dd | EE</td><td>0</td><td>0</td><td>1/2</td><td>1/4</td><td>1/4</td><td>1/4</td><td>1/4</td><td>0</td><td>1/2</td><td>0</td></tr>
<tr><td>SRSS</td><td>Bb~cc | Dd | Ee</td><td>0</td><td>1/4</td><td>0</td><td>1/8</td><td>1/8</td><td>1/4</td><td>0</td><td>1/4</td><td>0</td><td>1</td></tr>
<tr><td>SRSS</td><td>BB~cc | Dd | Ee</td><td>0</td><td>0</td><td>1/4</td><td>1/8</td><td>1/8</td><td>1/4</td><td>0</td><td>1/4</td><td>0</td><td>0</td></tr>
<tr><td>SRSS</td><td>Bb~cc | DD | Ee</td><td>1/2</td><td>1/4</td><td>0</td><td>1/8</td><td>1/8</td><td>0</td><td>1/4</td><td>1/4</td><td>0</td><td>0</td></tr>
<tr><td>SRSS</td><td>BB~cc | DD | Ee</td><td>0</td><td>0</td><td>1/4</td><td>1/8</td><td>1/8</td><td>0</td><td>1/4</td><td>1/4</td><td>0</td><td>0</td></tr>
<tr><td>SRSS</td><td>Bb~cc | Dd | EE</td><td>0</td><td>1/4</td><td>0</td><td>1/8</td><td>1/8</td><td>1/4</td><td>0</td><td>0</td><td>1/4</td><td>0</td></tr>
<tr><td>SRSS</td><td>BB~cc | Dd | EE</td><td>0</td><td>0</td><td>1/4</td><td>1/8</td><td>1/8</td><td>1/4</td><td>0</td><td>0</td><td>1/4</td><td>0</td></tr>
<tr><td>SRSS</td><td>Bb~cc | DD | EE</td><td>1/2</td><td>1/4</td><td>0</td><td>1/8</td><td>1/8</td><td>0</td><td>1/4</td><td>0</td><td>1/4</td><td>0</td></tr>
<tr><td>SRSS</td><td>BB~cc | DD | EE</td><td>0</td><td>0</td><td>1/4</td><td>1/8</td><td>1/8</td><td>0</td><td>1/4</td><td>0</td><td>1/4</td><td>0</td></tr>
<tr><td>SSRS</td><td>bb~cc | dd | ee</td><td>1/3</td><td>1/3</td><td>1/3</td><td>1/3</td><td>1/3</td><td>1/3</td><td>1/3</td><td>1</td><td>0</td><td>0</td></tr>
<tr><td>SSRS</td><td>bb~cc | dd | Ee</td><td>1/3</td><td>1/3</td><td>1/3</td><td>1/3</td><td>1/3</td><td>1/3</td><td>1/3</td><td>0</td><td>0</td><td>1</td></tr>
<tr><td>SSRS</td><td>bb~cc | dd | EE</td><td>1/3</td><td>1/3</td><td>1/3</td><td>1/3</td><td>1/3</td><td>1/3</td><td>1/3</td><td>0</td><td>1</td><td>0</td></tr>
<tr><td>SSSS</td><td>bb~cc | Dd | ee</td><td>0</td><td>1/6</td><td>1/6</td><td>1/6</td><td>1/6</td><td>1/3</td><td>0</td><td>1/2</td><td>0</td><td>0</td></tr>
<tr><td>SSSS</td><td>bb~cc | DD | ee</td><td>1/3</td><td>1/6</td><td>1/6</td><td>1/6</td><td>1/6</td><td>0</td><td>1/3</td><td>1/2</td><td>0</td><td>0</td></tr>
<tr><td>SSSS</td><td>bb~cc | Dd | Ee</td><td>0</td><td>1/6</td><td>1/6</td><td>1/6</td><td>1/6</td><td>1/3</td><td>0</td><td>0</td><td>0</td><td>1</td></tr>
<tr><td>SSSS</td><td>bb~cc | DD | Ee</td><td>1/3</td><td>1/6</td><td>1/6</td><td>1/6</td><td>1/6</td><td>0</td><td>1/3</td><td>0</td><td>0</td><td>0</td></tr>
<tr><td>SSSS</td><td>bb~cc | Dd | EE</td><td>0</td><td>1/6</td><td>1/6</td><td>1/6</td><td>1/6</td><td>1/3</td><td>0</td><td>0</td><td>1/2</td><td>0</td></tr>
<tr><td>SSSS</td><td>bb~cc | DD | EE</td><td>1/3</td><td>1/6</td><td>1/6</td><td>1/6</td><td>1/6</td><td>0</td><td>1/3</td><td>0</td><td>1/2</td><td>0</td></tr>
</table>

**Table S2** Resistance genotype distribution scenario in the Aegelsee *Daphnia magna* population, producing an expected resistotype distribution that fits best the observed one. We fix the "b" and the "D" alleles, and we distribute proportions in the remaining possible genotypes, within each resistotype, using C- and E-loci allele frequency observed in spring 2015. In resistotypes where it is not possible to fix the allele, we equally distribute genotype proportions among heterozygous genotypes or among homozygous genotypes for the alternative allele when this is the only possible genotype determining the resistotype. The genetic model described in Fig. S5 provided the list of possible genotypes and their corresponding resistotypes. This table corresponds to the "freq" table described in Supplementary Doc. S2 and Fig. S8.

| pheno | geno | | | Genotype distribution within resistotypes | |
|---|---|---|---|---|---|
| | | | | **bbDD scenario and C- and E-loci allele frequency inferred** | |
| RRRR | bb~Cc | dd | ee | x | $f_{RRRR}$(Cc)=f(Cc)/(f(Cc)+f(CC))=0.65 |
| RRRR | Bb~Cc | dd | ee | 0 | 0 |
| RRRR | BB~Cc | dd | ee | 0 | 0 |
| RRRR | bb~CC | dd | ee | x | $f_{RRRR}$(CC)=f(CC)/(f(Cc)+f(CC))=0.35 |
| RRRR | Bb~CC | dd | ee | 0 | 0 |
| RRRR | BB~CC | dd | ee | 0 | 0 |
| RRSR | bb~Cc | Dd | ee | 0 | 0 |
| RRSR | Bb~Cc | Dd | ee | 0 | 0 |
| RRSR | BB~Cc | Dd | ee | 0 | 0 |
| RRSR | bb~CC | Dd | ee | 0 | 0 |
| RRSR | Bb~CC | Dd | ee | 0 | 0 |
| RRSR | BB~CC | Dd | ee | 0 | 0 |
| RRSR | bb~Cc | DD | ee | x | $f_{RRSR}$(Cc)=f(Cc)/(f(Cc)+f(CC))=0.65 |
| RRSR | Bb~Cc | DD | ee | 0 | 0 |
| RRSR | BB~Cc | DD | ee | 0 | 0 |
| RRSR | bb~CC | DD | ee | x | $f_{RRSR}$(CC)=f(CC)/(f(Cc)+f(CC))=0.35 |
| RRSR | Bb~CC | DD | ee | 0 | 0 |
| RRSR | BB~CC | DD | ee | 0 | 0 |
| SRRR | Bb~cc | dd | ee | x | $f_{SRRR}$=1 |
| SRRR | BB~cc | dd | ee | 0 | 0 |
| SRSR | Bb~cc | Dd | ee | 0 | 0 |
| SRSR | BB~cc | Dd | ee | 0 | 0 |
| SRSR | Bb~cc | DD | ee | x | $f_{SRSR}$=1 |
| SRSR | BB~cc | DD | ee | 0 | 0 |
| RRRS | bb~Cc | dd | Ee | x | $f_{RRRS}$(Cc_Ee)=(f(Cc)*f(Ee))/((f(Cc)+f(CC))*(f(Ee)+f(EE)))=0.557 |
| RRRS | Bb~Cc | dd | Ee | 0 | 0 |
| RRRS | BB~Cc | dd | Ee | 0 | 0 |
| RRRS | bb~CC | dd | Ee | x | $f_{RRRS}$(CC_Ee)=(f(CC)*f(Ee))/((f(Cc)+f(CC))*(f(Ee)+f(EE)))=0.294 |
| RRRS | Bb~CC | dd | Ee | 0 | 0 |
| RRRS | BB~CC | dd | Ee | 0 | 0 |
| RRRS | bb~Cc | dd | EE | x | $f_{RRRS}$(Cc_EE)=(f(Cc)*f(EE))/((f(Cc)+f(CC))*(f(Ee)+f(EE)))=0.097 |
| RRRS | Bb~Cc | dd | EE | 0 | 0 |
| RRRS | BB~Cc | dd | EE | 0 | 0 |
| RRRS | bb~CC | dd | EE | x | $f_{RRRS}$(CC_EE)=(f(CC)*f(EE))/((f(Cc)+f(CC))*(f(Ee)+f(EE)))=0.052 |
| RRRS | Bb~CC | dd | EE | 0 | 0 |
| RRRS | BB~CC | dd | EE | 0 | 0 |
| RRSS | bb~Cc | Dd | Ee | 0 | 0 |
| RRSS | Bb~Cc | Dd | Ee | 0 | 0 |
| RRSS | BB~Cc | Dd | Ee | 0 | 0 |
| RRSS | bb~CC | Dd | Ee | 0 | 0 |
| RRSS | Bb~CC | Dd | Ee | 0 | 0 |
| RRSS | BB~CC | Dd | Ee | 0 | 0 |
| RRSS | bb~Cc | DD | Ee | x | $f_{RRSS}$(Cc_Ee)=(f(Cc)*f(Ee))/((f(Cc)+f(CC))*(f(Ee)+f(EE)))=0.557 |
| RRSS | Bb~Cc | DD | Ee | 0 | 0 |
| RRSS | BB~Cc | DD | Ee | 0 | 0 |
| RRSS | bb~CC | DD | Ee | x | $f_{RRSS}$(CC_Ee)=(f(CC)*f(Ee))/((f(Cc)+f(CC))*(f(Ee)+f(EE)))=0.294 |
| RRSS | Bb~CC | DD | Ee | 0 | 0 |
| RRSS | BB~CC | DD | Ee | 0 | 0 |
| RRSS | bb~Cc | Dd | EE | 0 | 0 |
| RRSS | Bb~Cc | Dd | EE | 0 | 0 |
| RRSS | BB~Cc | Dd | EE | 0 | 0 |
| RRSS | bb~CC | Dd | EE | 0 | 0 |
| RRSS | Bb~CC | Dd | EE | 0 | 0 |
| RRSS | BB~CC | Dd | EE | 0 | 0 |
| RRSS | bb~Cc | DD | EE | x | $f_{RRSS}$(Cc_EE)=(f(Cc)*f(EE))/((f(Cc)+f(CC))*(f(Ee)+f(EE)))=0.097 |
| RRSS | Bb~Cc | DD | EE | 0 | 0 |
| RRSS | BB~Cc | DD | EE | 0 | 0 |
| RRSS | bb~CC | DD | EE | x | $f_{RRSS}$(CC_EE)=(f(CC)*f(EE))/((f(Cc)+f(CC))*(f(Ee)+f(EE)))=0.052 |
| RRSS | Bb~CC | DD | EE | 0 | 0 |
| RRSS | BB~CC | DD | EE | 0 | 0 |
| SRRS | Bb~cc | dd | Ee | x | $f_{SRRS}$(Ee)=f(Ee)/(f(Ee)+f(EE))=0.85 |
| SRRS | BB~cc | dd | Ee | 0 | 0 |
| SRRS | Bb~cc | dd | EE | x | $f_{SRRS}$(EE)=f(EE)/(f(Ee)+f(EE))=0.15 |
| SRRS | BB~cc | dd | EE | 0 | 0 |
| SRSS | Bb~cc | Dd | Ee | 0 | 0 |
| SRSS | BB~cc | Dd | Ee | 0 | 0 |
| SRSS | Bb~cc | DD | Ee | x | $f_{SRSS}$(Ee)=f(Ee)/(f(Ee)+f(EE))=0.85 |
| SRSS | BB~cc | DD | Ee | 0 | 0 |
| SRSS | Bb~cc | Dd | EE | 0 | 0 |
| SRSS | BB~cc | Dd | EE | 0 | 0 |
| SRSS | Bb~cc | DD | EE | x | $f_{SRSS}$(EE)=f(EE)/(f(Ee)+f(EE))=0.15 |
| SRSS | BB~cc | DD | EE | 0 | 0 |
| SSRS | bb~cc | dd | ee | x | $f_{SSRS}$(ee)=f(ee)=0.55 |
| SSRS | bb~cc | dd | Ee | x | $f_{SSRS}$(Ee)=f(Ee)=0.38 |
| SSRS | bb~cc | dd | EE | x | $f_{SSRS}$(EE)=f(EE)=0.07 |
| SSSS | bb~cc | Dd | ee | 0 | 0 |
| SSSS | bb~cc | DD | ee | x | $f_{SSSS}$(ee)=f(ee)=0.55 |
| SSSS | bb~cc | Dd | Ee | 0 | 0 |
| SSSS | bb~cc | DD | Ee | x | $f_{SSSS}$(Ee)=f(Ee)=0.38 |
| SSSS | bb~cc | Dd | EE | 0 | 0 |
| SSSS | bb~cc | DD | EE | x | $f_{SSSS}$(EE)=f(EE)=0.07 |

**Table S3** Participation of the different resistotypes to sexual reproduction in the host population. To assess whether sexual reproduction was biased towards some resistotypes, we collected in August 2020 *Daphnia magna* samples and quantified the resistotype distribution of females carrying resting stages, of males and of a random sample of females. Since males cannot be cloned, we cannot produce replicates to assess their resistotype, resulting in higher uncertainty of their resistotypes, in particular for the less easy scorable P15 and P21 *Pasteuria ramosa* isolates.

| | Resistotype | | | | | | | | |
|---|---|---|---|---|---|---|---|---|---|
| | RRRRR | RRRRS | RRSRR | RRSRS | RRSSS | RSRSR | RSSRS | SRSRS | SSSRS |
| Field sample 5. August 2020: | 0 | 1 | 7 | 70 | 4 | 0 | 1 | 1 | 0 |
| Field sample 25. August 2020: | 2 | 2 | 1 | 72 | 1 | 0 | 1 | 0 | 1 |
| Males (pooled both dates): | 4 | 10 | 6 | 73 | 1 | 1 | 2 | 0 | 0 |
| Ephippial mothers (pooled both dates): | 0 | 0 | 0 | 79 | 0 | 0 | 0 | 0 | 0 |

**Methods**

***Hatching modelling***

**Document S1** peas implementation of the genetic model of resistance in the *Daphnia magna* – *Pasteuria ramosa* system.

### Genetic model for resistance in the Aegelsee

implemented in peas R package

#### BCDE genetic model

#### 1. Install the package

```
#install.packages("devtools")
#devtools::install_github("JanEngelstaedter/peas", build_vignettes = TRUE)

library(peas)
```

#### 2. Set up the genetic model

##### 2.1 Defining the genetic system

```
r2<-(1-exp(-2*23.1/100))/2 ## recombination rate between the B and C loci (Metzger et al. 201
6)

# 4 loci with 2 alleles each, BC clustered togheter (Metzger et al. 2016).
BCDE <- newGenopheno(nloci = 2,
           alleleNames = list(c("b", "B"), c("c", "C")),
           rec = r2)
BCDE <- addLinkageGroup(BCDE, alleleNames = list(c("d", "D")))
BCDE <- addLinkageGroup(BCDE, alleleNames = list(c("e", "E")))
```

##### 2.2 Set genotypes and their corresponding phenotypes : THE GENETIC MODEL

```
BCDE <- setPhenotypes(BCDE, "S/R", "_~_|_|_","SSRR") # default all rececive --> SSRR
# (we don't take the epistasis relation into account, this will be the last line of the model)

BCDE <- setPhenotypes(BCDE, "S/R", "B_~_|_|_","SRRR") # B --> R to C19

BCDE <- setPhenotypes(BCDE, "S/R", "B_~_|D_|_","SRSR") # D --> S to P15
# (Bento et al. 2020)
BCDE <- setPhenotypes(BCDE, "S/R", "B_~_|_|E_","SRRS") # E --> S to P20
BCDE <- setPhenotypes(BCDE, "S/R", "B_~_|D_|E_","SRSS") #


BCDE <- setPhenotypes(BCDE, "S/R", "_~C_|_|_","RRRR") # C hides B and --> R to C1 and C1
```

```
9

BCDE <- setPhenotypes(BCDE, "S/R", "__~C_ | D_ | __","RRSR") #
BCDE <- setPhenotypes(BCDE, "S/R", "__~C_ | __ | E_","RRRS") #
BCDE <- setPhenotypes(BCDE, "S/R", "__~C_ | D_ | E_","RRSS") #

# b/c-E epistasis #
# "bbcc" genotype induces S to P20, regardless of genotype at E-locus

BCDE <- setPhenotypes(BCDE, "S/R", "bb~cc | D_ | __","SSSS") #
BCDE <- setPhenotypes(BCDE, "S/R", "bb~cc | dd | __","SSRS") #
```

*Summary of model*

```
BCDE

## Genetic system comprising 3 linkage groups:
##   Linkage group 1: autosomal, 2 loci with recombination rate 0.1849888
##     Alleles at locus 1: b, B
##     Alleles at locus 2: c, C
##   Linkage group 2: autosomal, 1 locus
##     Alleles at locus 1: d, D
##   Linkage group 3: autosomal, 1 locus
##     Alleles at locus 1: e, E
## Phenotypes defined for the following traits:
##   S/R (trait values: SSRS, SRRR, RRRR, SSSS, SRSR, RRSR, SRRS, RRRS, SRSS, RRSS)
```

*List of all possible genotype combinations and corresponding phenotypes*

```
BCDEallgeno<-getPhenotypes(BCDE)
nrow(BCDEallgeno) # 81 possible genotypes

## [1] 81

nrow(unique(BCDEallgeno)) # 10 possible resistotypes

## [1] 10

BCDEallgeno$geno<-row.names(BCDEallgeno) # add "geno" column
BCDEallgeno<-BCDEallgeno[order(BCDEallgeno$`S/R`),] # sort resistotypes
# rename "S/R" column as "pheno"
library(tidyverse)
BCDEallgeno<-rename(BCDEallgeno, pheno=`S/R`)
# export in xl file
library(xlsx)
write.xlsx(BCDEallgeno,"BCDEallgeno.xlsx")
```

### 3. Predict crosses

```
# all possible genotype crossings and their
# expected F1 genotype and phenotype segregation


cross<-matrix(list(), nrow=nrow(BCDEallgeno),
       ncol=nrow(BCDEallgeno), byrow=T)



for (i in 1:nrow(BCDEallgeno)) {
```

```
 for (j in 1:nrow(BCDEallgeno)) {
  cross[[i,j]]<-predictCross(BCDE, BCDEallgeno$geno[i],BCDEallgeno$geno[j])
  # we add the corresponding resistotypes to the genotype output
  cross[[i,j]]$genotypes$SRtrait <- getPhenotypes(BCDE, equivalent = "none")[row.names(cr
oss[[i,j]]$genotypes), ]
 }
}

# large matrix of 81*81=6561 elements
# example: cross between genotype#1 and genotype#2
BCDEallgeno[1,]

##               pheno          geno
## bb~Cc | dd | ee  RRRR bb~Cc | dd | ee

BCDEallgeno[2,]

##               pheno          geno
## Bb~Cc | dd | ee  RRRR Bb~Cc | dd | ee

cross[[1,2]]

## $genotypes
##                 fraction SRtrait
## bB~CC | dd | ee 0.20375279    RRRR
## bb~CC | dd | ee 0.04624721    RRRR
## bB~Cc | dd | ee 0.25000000    RRRR
## bb~Cc | dd | ee 0.25000000    RRRR
## bB~cc | dd | ee 0.04624721    SRRR
## bb~cc | dd | ee 0.20375279    SSRS
##
## $phenotypes
##    S/R   fraction
## 1 RRRR 0.75000000
## 2 SRRR 0.04624721
## 3 SSRS 0.20375279

# find a genotype
match("Bb~cc | dd | EE", BCDEallgeno$geno) # position 59

## [1] 59
```

**Doc. S2 and Fig. S8**: Theoretical resistotype frequency resulting from ephippia hatching in the Aegelsee.

Using the genetic model of resistance in the *Daphnia magna* - *Pasteuria ramosa* system, we calculate theoretical resistotype (resistance phenotype) frequency resulting from hatching of *D. magna* resting stages laid throughout the active season.

*Note*: we use four-letter resistotype because the genetic model of resistance includes resistance to the four *P. ramosa* isolates C1, C19, P15 and P20.

**Document S2** calculations of the theoretical resistance phenotype frequency resulting from ephippia hatching in the Aegelsee.

**prop data frame**: observed resistotype proportion in the study population at each sampling date.

**prop**

| | pheno-1 (RRRR) | ... | pheno-**k** | ... | pheno-n (SSSS) |
|---|---|---|---|---|---|
| date-1 | $p_{1,1}$ | ... | $p_{1,k}$ | ... | $p_{1,n}$ |
| date-2 | $p_{2,1}$ | ... | $p_{2,k}$ | ... | $p_{2,n}$ |
| date-3 | $p_{3,1}$ | ... | $p_{3,k}$ | ... | $p_{3,n}$ |
| ... | ... | ... | ... | ... | ... |
| date-**i** | $p_{i,1}$ | ... | $\mathbf{p_{i,k}}$ | ... | $p_{i,n}$ |
| ... | ... | ... | ... | ... | ... |
| date-d | $p_{d,1}$ | ... | $p_{d,k}$ | ... | $p_{d,n}$ |

**freq data frame**: fraction of each possible genotype within each phenotype. We use the list of possible genotypes and their corresponding phenotypes given by the genetic model of resistance implemented in the "peas" R-package. This list is given by the "getPhenotypes" function in the "peas" R-package (Doc. S1). The fraction of each possible genotype within each phenotype (fq) is implemented by the user. In Figs. S6 and S7, Tables S1 and S2 we test different scenarios of genotype distribution within the resistotypes.

**freq**

| pheno | geno | fq |
|---|---|---|
| pheno-1 (RRRR) | geno-1 | $f_{1,1}$ |
| pheno-1 (RRRR) | geno-2 | $f_{2,1}$ |
| pheno-1 (RRRR) | geno-3 | $f_{3,1}$ |
| pheno-2 | geno-4 | $f_{4,2}$ |
| pheno-2 | geno-5 | $f_{5,2}$ |
| pheno-2 | geno-6 | $f_{6,2}$ |
| ... | ... | ... |
| pheno-**k** | ... | ... |
| pheno-**k** | ... | ... |
| pheno-**k** | ... | ... |
| pheno-**k** | geno-**j** | $\mathbf{f_{j,k}}$ |
| ... | ... | ... |
| pheno-n (SSSS) | geno-g | $f_{g,n}$ |

We then calculate the proportion of each genotype at each sampling date:

$$d_{j,i} = f_{j,k} * p_{i,k}$$

with

$\mathbf{d_{j,i}}$ : proportion of the j-genotype at the i-date.

$\mathbf{f_{j,k}}$ from the **freq** data frame: proportion of the j-genotype within the k-phenotype. This is implemented by the user.

$\mathbf{p_{i,k}}$ from the **prop** data frame: proportion of the k-phenotype at the i-date. Input from sampled data in the study population.

**freq**

| pheno | geno | fq | date-1 | date-2 | ... | date-**i** | ... | date-d |
|---|---|---|---|---|---|---|---|---|
| pheno-1 | geno-1 | $f_{1,1}$ | $d_{1,1}$ | $d_{1,2}$ | ... | $d_{1,i}$ | ... | $d_{1,d}$ |
| pheno-1 | geno-2 | $f_{2,1}$ | $d_{2,1}$ | $d_{2,2}$ | ... | $d_{2,i}$ | ... | $d_{2,d}$ |
| pheno-1 | geno-3 | $f_{3,1}$ | $d_{3,1}$ | $d_{3,2}$ | ... | $d_{3,i}$ | ... | $d_{3,d}$ |
| pheno-2 | geno-4 | $f_{4,2}$ | $d_{4,1}$ | $d_{4,2}$ | ... | $d_{4,i}$ | ... | $d_{4,d}$ |
| pheno-2 | geno-5 | $f_{5,2}$ | $d_{5,1}$ | $d_{5,2}$ | ... | $d_{5,i}$ | ... | $d_{5,d}$ |
| pheno-2 | geno-6 | $f_{6,2}$ | $d_{6,1}$ | $d_{6,2}$ | ... | $d_{6,i}$ | ... | $d_{6,d}$ |
| ... | ... | ... | ... | ... | ... | ... | ... | ... |
| pheno-**k** | ... | ... | ... | ... | ... | ... | ... | ... |
| pheno-**k** | ... | ... | ... | ... | ... | ... | ... | ... |
| pheno-**k** | ... | ... | ... | ... | ... | ... | ... | ... |
| pheno-**k** | geno-**j** | $f_{j,k}$ | $d_{j,1}$ | $d_{j,2}$ | ... | $\mathbf{d_{j,i}}$ | ... | $d_{j,d}$ |
| ... | ... | ... | ... | ... | ... | ... | ... | ... |
| pheno-n | geno-g | $f_{g,n}$ | $d_{g,1}$ | $d_{g,2}$ | ... | $d_{g,i}$ | ... | $d_{g,d}$ |

**di matrix**: fraction of all possible genotype crossings at the i-date considering random mating and equal contribution of all individuals to sexual reproduction.

$$di = \begin{pmatrix} d_{1,i} \\ d_{2,i} \\ \vdots \\ d_{j,i} \\ \vdots \\ d_{g,i} \end{pmatrix} * t \begin{pmatrix} d_{1,i} \\ d_{2,i} \\ \vdots \\ d_{j,i} \\ \vdots \\ d_{g,i} \end{pmatrix} = \begin{pmatrix} di_{1,1} & di_{1,2} & \cdots & di_{1,m} & \cdots & di_{1,g} \\ di_{2,1} & di_{2,2} & \cdots & di_{2,m} & \cdots & di_{2,g} \\ \vdots & \vdots & \ddots & \vdots & \ddots & \vdots \\ di_{l,1} & di_{l,2} & \cdots & di_{l,m} & \cdots & di_{l,g} \\ \vdots & \vdots & \ddots & \vdots & \ddots & \vdots \\ di_{g,1} & di_{g,2} & \cdots & di_{g,m} & \cdots & di_{g,g} \end{pmatrix}$$

with

$\mathbf{di_{l,m}}$: proportion of mating events between the l-genotype and m-genotype at the i-date.

The sum of the elements of the di matrix is equal to 1.

**cross matrix**

$$\begin{pmatrix} c_{1,1} & c_{1,2} & \cdots & c_{1,m} & \cdots & c_{1,g} \\ c_{2,1} & c_{2,2} & \cdots & c_{2,m} & \cdots & c_{2,g} \\ \vdots & \vdots & \ddots & \vdots & \ddots & \vdots \\ c_{l,1} & c_{l,2} & \cdots & c_{l,m} & \cdots & c_{l,g} \\ \vdots & \vdots & \ddots & \vdots & \ddots & \vdots \\ c_{g,1} & c_{g,2} & \cdots & c_{g,m} & \cdots & c_{g,g} \end{pmatrix}$$

$\mathbf{c_{l,m}}$: data frame containing predicted genotypic and phenotypic crossing results between the l- and the m-genotype. This was calculated using the “predictCross” function in the “peas” R-package. See implementation of the genetic model of resistance in Doc. S1.

Example:

$c_{1,2}$: crossing result between genotype #1 and genotype #2

```
BCDEallgeno[1,]
```

```
##            pheno         geno
## bb~Cc | dd | ee  RRRR bb~Cc | dd | ee

BCDEallgeno[2,]

##            pheno         geno
## Bb~Cc | dd | ee  RRRR Bb~Cc | dd | ee

cross[[1,2]]

## $genotypes
##              fraction SRtrait
## bB~CC | dd | ee 0.20375279    RRRR
## bb~CC | dd | ee 0.04624721    RRRR
## bB~Cc | dd | ee 0.25000000    RRRR
## bb~Cc | dd | ee 0.25000000    RRRR
## bB~cc | dd | ee 0.04624721    SRRR
## bb~cc | dd | ee 0.20375279    SSRS
##
## $phenotypes
##    S/R   fraction
## 1 RRRR 0.75000000
## 2 SRRR 0.04624721
## 3 SSRS 0.20375279
```

**$c_{l,m}$$genotypes**

| | fraction | SRtrait |
|---|---|---|
| geno-a | $\alpha_a$ | pheno-a |
| geno-b | $\alpha_b$ | pheno-a |
| geno-c | $\alpha_c$ | pheno-b |
| ... | ... | ... |
| geno-x | $\alpha_x$ | pheno-y |

With y ≤ x as there can be several genotypes underlying one phenotype (see example above)

**$c_{l,m}$$phenotypes**

| | S/R | fraction |
|---|---|---|
| 1 | pheno-a | $\beta_a$ |
| 2 | pheno-b | $\beta_b$ |
| ... | ... | ... |
| z | pheno-y | $\beta_y$ |

**prophatch data frame**: final data frame with expected resistotype proportions at each sampling date.

**prophatch**

| | pheno-1 (RRRR) | ... | pheno-**k** | ... | pheno-n (SSSS) |
|---|---|---|---|---|---|
| date-1 | $h_{1,1}$ | ... | $h_{1,k}$ | ... | $h_{1,n}$ |
| date-2 | $h_{2,1}$ | ... | $h_{2,k}$ | ... | $h_{2,n}$ |
| date-3 | $h_{3,1}$ | ... | $h_{3,k}$ | ... | $h_{3,n}$ |
| ... | ... | ... | ... | ... | ... |
| date-**i** | $h_{i,1}$ | ... | $\mathbf{h_{i,k}}$ | ... | $h_{i,n}$ |
| ... | ... | ... | ... | ... | ... |
| date-d | $h_{d,1}$ | ... | $h_{d,k}$ | ... | $h_{d,n}$ |

$\mathbf{h_{i,k}}$: expected frequency of the k-resistotype at the i-date.

$$h_{i,k} = \sum_{m=0}^{j} \sum_{l=0}^{j} c_{l,m}\$phenotypes\$\beta_k * di_{l,m}$$

The same calculation can be done with the genotypes fractions (α) instead of the phenotypes fractions (β).

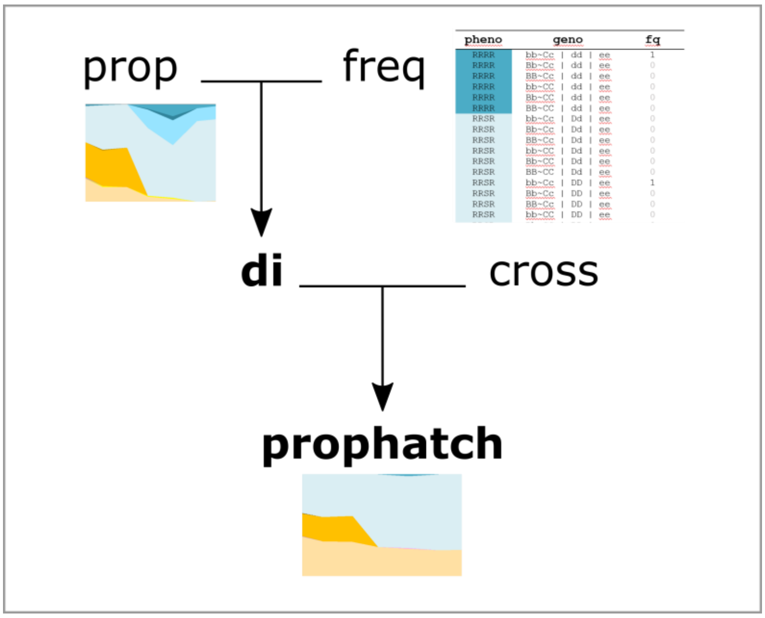


**Figure S8** Summary illustration of Doc. S2: calculation of theoretical resistotype (resistance phenotype) frequency resulting from resting stages hatching in the Aegelsee *Daphnia magna* population. The theoretical resistotype frequency over time is calculated using the genetic model of resistance in the *D. magna* - *Pasteuria ramosa* system and the observed resistotype frequency over time in the *D. magna* population. Input elements are written in regular font style, output elements are written in bold. **Prop** data frame: observed resistotype proportion in the study population at each sampling date. **Freq** data frame: fraction of each possible genotype within each phenotype. We use the list of possible genotypes and their corresponding phenotypes given by the genetic model of resistance implemented in the "peas" R-package. **Di** matrix: calculated from "prop" and "freq": fraction of all possible genotype crossings at the i-date considering random mating in the *D. magna* population. **Cross** matrix: predicted genotypic and phenotypic crossing results from all possible genotype crossings. Each element of the matrix is a data frame containing predicted genotypic and phenotypic crossing results of one genotype crossing. **Prophatch** data frame: calculated from "di" and "cross": final data frame with expected resistotype proportions resulting from sexual reproduction at each sampling date.
